# Benchmarking common preprocessing strategies in early childhood functional connectivity MRI

**DOI:** 10.1101/2020.10.27.358192

**Authors:** Kirk Graff, Ryann Tansey, Amanda Ip, Christiane Rohr, Dennis Dimond, Deborah Dewey, Signe Bray

**Author notes:** Corresponding author: Kirk Graff, Alberta Children’s Hospital Research Institute B4-290, 28 Oki Drive NW, Calgary, AB, T3B 6A8, Canada.

## Abstract

Functional connectivity magnetic resonance imaging (FC-MRI) has been widely used to investigate neurodevelopment. However, FC-MRI is vulnerable to head motion, which is associated with age and distorts FC estimates. Numerous preprocessing strategies have been developed to mitigate confounds, each with advantages and drawbacks. Preprocessing strategies for FC-MRI have typically been validated and compared using resting state data from adults. However, FC-MRI in young children presents a unique challenge due to relatively high head motion and a growing use of passive viewing paradigms to mitigate motion. This highlights a need to compare processing choices in pediatric samples. To this end, we leveraged longitudinal, passive viewing fMRI data collected from 4 to 8-year-old children. We systematically investigated combinations of widely used and debated preprocessing strategies, namely global signal regression, volume censoring, ICA-AROMA, and bandpass filtering. We implemented commonly used metrics of noise removal (i.e. quality control-functional connectivity), metrics sensitive to individual differences (i.e. connectome fingerprinting), and, because data was collected during a passive viewing task, we also assessed the impact on stimulus-evoked responses (i.e. intersubject correlations; ISC). We found that the most efficacious pipeline included censoring, global signal regression, bandpass filtering, and head motion parameter regression. Despite the drawbacks of noise-mitigation steps, our findings show benefits for both noise removal and information retention in a high-motion early childhood sample.

**Highlights:**

- We evaluated 27 preprocessing pipelines in passive viewing data from young children
- Pipelines were evaluated on noise-removed and information retained
- Pipelines that included censoring and GSR outperformed alternatives across benchmarks
- For high-motion scans, preprocessing choices substantially alter connectomes

## 1. Introduction

Functional connectivity magnetic resonance imaging (FC-MRI) enables investigation of functional networks in the brain. While FC-MRI identifies consistent FC patterns across individuals (Fox et al., 2005; Damoiseaux et al., 2006), it is highly sensitive to artifacts from physiological sources, such as heart rate and respiration, and head motion (Power et al., 2012; Satterthwaite et al., 2012; Van Dijk et al., 2012). This is particularly problematic in developmental neuroimaging studies, which face the challenge of age-correlated head motion (Power et al., 2012; Fair et al., 2012). To reduce head motion and increase compliance, FC-MRI studies in children are increasingly conducted using passive viewing tasks such as movies (Rohr et al., 2017; Greene et al., 2018). However, head motion noise remains a major concern, and preprocessing choices are therefore important in studies of young children. There is currently no gold-standard preprocessing pipeline in the literature, for pediatric studies or otherwise, and noise mitigation steps involve tradeoffs and remain widely debated (Murphy and Fox, 2017; Satterthwaite et al., 2019). Common strategies include regression of motion estimates, independent component analysis (ICA)-based approaches, regression of the global signal, censoring volumes of high framewise displacement, and temporal filtering.

Regressing out motion estimates remains one of the most common denoising approaches (Satterthwaite et al., 2019). However, based on pipeline benchmarks, Ciric et al. (2017) suggest that this approach is insufficient on its own and leads to concerns about losses in degrees of freedom. An alternative approach is to remove effects of head motion through ICA-based approaches, such as ICA-AROMA (Pruim et al., 2015), which decomposes data into components that reflect either brain activity or structured noise. ICA-AROMA automatically classifies components as noise by both temporal features (high frequency content, correlation with realignment parameters) and spatial features (near CSF or the edge of the brain) (Pruim et al., 2015). These structured noise components can then be regressed from the data (Thomas et al., 2002).

Global signal regression (GSR) is an often used but widely debated preprocessing step (Murphy and Fox, 2017; Chai et al., 2012; Gotts et al., 2013). GSR is a simple and arguably effective (Ciric et al., 2017; Parkes et al., 2018) denoising technique, improving the specificity of positive correlations and showing results that are more consistent with anatomical connectivity (Fox et al., 2009). However, the use of GSR tends to increase the apparent strength of short-range connections while decreasing that of long-range connections (Saad et al., 2012; Ciric et al., 2017; Parkes et al., 2018), decreases positive BOLD responses, and creates anti-correlations which may not exist (Aguirre et al., 1998). The global signal has been shown to resemble established networks and to be significantly related to life outcomes and psychological function, suggesting it is rich in information on cognition and behavior (Li et al., 2019), and that by extension GSR should be used with caution as it removes information specific to the individual.

To mitigate the effect of specific motion contaminated volumes, it has become common to either remove, or interpolate over, specific time points by censoring or ‘scrubbing’ (Power et al., 2012). Despite its apparent efficacy in removing noise (Ciric et al., 2017), censoring has concerns, such as disrupting temporal autocorrelations, and leaving variable amounts of scan data between participants. Further, even if the number of censored volumes is matched, not all censored volumes are equally rich in information (Power et al., 2015). Censoring regimes vary widely across studies (Satterthwaite et al., 2013; Power et al., 2012; Power et al., 2014) and it is a challenge to determine which level of censoring optimizes the recovery of individually specific FC information.

Another preprocessing strategy is temporal filtering. fMRI signals at both very low frequencies (under 0.01 Hz) and high frequencies (above 0.1 Hz) are often filtered out to remove noise (Satterthwaite et al., 2013). However, filtering above 0.1 Hz may also be removing connectivity information (Niazy et al., 2011), or artificially increasing correlations by introducing sample dependence (Davey et al., 2013), leading to concern on the appropriateness of a bandpass filter compared to a highpass filter (Satterthwaite et al., 2019).

In response to the proliferation of preprocessing approaches (Carp, 2012), several studies have compared the effectiveness of different pipelines (Churchill et al., 2017; Ciric et al., 2017; Parkes et al., 2018). However, to our knowledge, no studies have systematically compared preprocessing pipelines in pediatric samples nor in studies that have used passive viewing tasks, which is important for several reasons. Passive viewing tasks alter FC relative to rest (Bray et al., 2015; Vanderwal et al., 2017), and have been shown to provide more stable FC estimates (Wang et al., 2017), which in turn may result in enhanced ability to identify brain-behavior correlations (Vanderwal et al., 2019). It remains unknown to what extent these FC changes are impacted by preprocessing choices. Young children typically have higher head motion than adults (Dosenbach et al., 2017), however, their smaller head size could alter the impact of rotational motion. Similarly, Bolton et al. (2020) found that in-scanner movement patterns are related to anthropometric and cognitive factors, suggesting it may be inappropriate to treat motion in children the same as motion in adults. Children also have faster respiratory rates and heart rates than adults (Fleming et al., 2011), which likely alters the physiological noise present in scans, and which is not accounted for with benchmarks that only account for head motion, such as the most commonly used previous metric, quality control – functional connectivity (QC-FC) (Satterthwaite et al., 2013; Power et al., 2012). At the same time, there are fundamental differences in the brain structure of children relative to adults (Grayson and Fair, 2017), and since GSR shows distance dependent effects, this step warrants investigation in children specifically.

Preprocessing steps that aggressively remove signal related to motion involve tradeoffs. Ciric et al. (2017) speculated that de-noising methods could improve QC-FC metrics and improve reliability by removing both signal and noise, but in the process lose sensitivity to individual differences of interest. These individual differences may include FC correlates of fluid intelligence (Finn et al., 2015), developmental changes (Kaufmann et al., 2017), or symptom load (Byrge and Kennedy, 2020), which may themselves be correlated with motion (Power et al., 2015). A complementary metric that provides information about the individual-specific information remaining following preprocessing is functional connectome fingerprinting (Finn et al., 2015; subsequently referred to simply as “fingerprinting”), which aims to match scans from the same participant through FC estimates. Preprocessing strategies that either fail to remove noise, or remove too much signal of interest, will have poor individual identifiability.

Additionally, previous benchmarking studies using resting-state FC-MRI scans have not assessed the effect of noise mitigation steps on stimulus evoked functional responses. Such responses can be analyzed using intersubject correlation (ISC; Hasson et al., 2004), the temporal signal correlation of the same brain region between individuals during passive viewing of a video stimulus. Although distinct from FC, which is the main focus of this paper, this analysis provides information about how stimulus evoked responses are enhanced or suppressed by noise mitigation steps, thereby providing a more comprehensive picture of how these steps impact functional time series.

The present study systematically compares common noise-mitigation steps in data collected from young children, using QC-FC, fingerprinting, and ISC. The aim of this study is to support researchers conducting FC-MRI in young samples who use naturalistic imaging methods to consider the tradeoffs and effectiveness of several common preprocessing steps.

## 2. Methods

### 2.1. Participants

Data were from a study of early childhood brain development that included both single timepoint and longitudinal measurements (Dimond et al., 2020a; Dimond et al., 2020b; Rohr et al., 2019; Rohr et al., 2017). Participants were recruited from the local community through advertisements and through existing databases. All procedures were approved by the University of Calgary Conjoint Health Research Ethics Board. Parents provided informed consent and children provided assent to participate. Participants were children between 4 and 7 years of age at baseline without any major health concerns and were excluded if they had full-scale IQ more than 2 standard deviations below the mean (100), a history of neurodevelopment or psychiatric disorders, or any neurological diagnoses. 168 participants were scanned at baseline. Of those, 139 provided usable baseline data (i.e. free of excessive head motion), and of those 59 (15 male) produced usable 12-month follow-up data. From the sample of participants with both baseline and follow-up data, participants were included if after volume-wise censoring of the fMRI data (described in more detail below), both scans retained at least 11 minutes of uncensored (retained) data. 56 of the 59 children (14 male) reached this threshold, contributing 112 scans.

### 2.2. Data collection

Children were scanned at the Alberta Children’s Hospital. During fMRI scans a passive-viewing task was used, where participants watched clips from a children’s television show (Elmo’s World) for 1100 s. Prior to scans, children underwent a practice scan in an MRI simulator during which they watched the video that would be used during the fMRI scan and practiced staying still. Data was acquired with a 3T GE MR750 w (Waukesha, WI) scanner using a 32-channel head coil. An anatomical scan was acquired using a T1w 3D BRAVO sequence (TR = 6.764 ms, TE = 2.908 ms, FA = 10°, voxel size 0.8×0.8×0.8 mm^3^). fMRI scans were collected using a gradient-echo EPI sequence (TR = 2.5 s, TE = 30 ms, FA = 70°, voxel size 3.5×3.5×3.5 mm^3^).

### 2.3. Higher-vs. lower-motion subgroups

Preprocessing pipelines were compared using three categories of metrics using the whole sample of 112 scans. To further assess whether preprocessing pipelines were particularly advantageous for scans with relatively lower or higher amounts of head motion, we median-split the 112 scans into two groups of 56 scans, then re-calculated QC-FC metrics and ISC metrics described below.

### 2.4. Data and code availability

Raw data used in this study are not publicly available, in agreement with the terms of consent given by participants and the Conjoint Health and Research Ethics Board at the University of Calgary. Benchmarking results (e.g. QC-FC, fingerprinting, ISC values) for each pipeline, or other pertinent data, will be made available upon request. Software packages mentioned (Nipype, FSL, AFNI, ANTs) provided on their respective websites. Python scripts available at https://github.com/BrayNeuroimagingLab/BNL_open/tree/main/fMRI_preprocessing.

### 2.5. Basic preprocessing steps common across pipelines

The following preprocessing steps were run on all scans prior to the custom elements of pipelines, described below. All preprocessing was carried out with custom Python scripts integrating Nipype functionality (version 1.1.5; Gorgolewski et al., 2011) using FSL version 6.0.0 (Smith et al., 2004), ANTs version 3.0.0.0 (Avants et al., 2011), and AFNI version 18.3.03 (Cox, 1996). Scans from the same individual were preprocessed separately. Structural (T1w) images were preprocessed using ANTs. This included bias field correction, brain extraction, and tissue segmentation. Tissue segments were eroded using AFNI (CSF eroded twice, WM eroded 7 times).

Basic preprocessing for EPI data was as follows, largely following the procedure described by Ciric et al. (2018): a) FSL *MCFLIRT* to estimate head motion parameters (HMPs) and average relative framewise displacement (Jenkinson et al., 2002). b) FSL *slicetimer* for slice time correction. c) FSL *MCFLIRT* for rigid body realignment. We followed the recommendation in Power et al. (2017) to estimate HMP on raw data but carry out slice time correction prior to rigid body realignment, necessitating two FSL *MCFLIRT* steps. d) FSL *BET* to skull-strip EPI images (Smith, 2002). e) ANTs *Registration* (Avants et al., 2011) to generate a transformation matrix to warp the EPI image to a study-specific EPI template. This template was produced based on the procedure described by Huang et al. (2010). Specifically, a 3D EPI reference image was taken from each fMRI scan, chosen as a volume of low motion approximately in the middle of the scan. These references were warped to MNI space, then averaged together and smoothed to create the final study-specific template. f) FSL *FLIRT* boundary-based registration (Jenkinson et al., 2002) was used to generate a transformation matrix to warp the EPI image to the T1w image, then the inverse transformation matrix was used to warp tissue segmentations to functional image space. All preprocessing and confound mitigation steps were carried out in native space. g) A linear regression to remove the mean and linear and quadratic trends from each voxel was conducted. For pipelines that include censoring, time points marked for censoring were excluded from the model to calculate linear and quadratic trends. In pipelines without censoring, all timepoints were included.

### 2.6. Preprocessing pipelines tested

After first-stage preprocessing, pipelines varied systematically in whether they used GSR, censoring, and ICA-AROMA, as shown in Table 1, creating pipelines A1 through A8.

**Table 1:** Preprocessing pipelines tested to compare effects of ICA-AROMA, global signal regression and censoring.

| Pipeline | Primary motion artifact removal | Global signal regression (GSR) | Censoring |
| --- | --- | --- | --- |
| A1 | Regress head motion parameters (HMPs) |  |  |
| A2 |  | Yes |  |
| A3 |  |  | Yes |
| A4 |  | Yes | Yes |
| A5 | ICA-AROMA |  |  |
| A6 |  | Yes |  |
| A7 |  |  | Yes |
| A8 |  | Yes | Yes |

### 2.7. Additional preprocessing

#### 2.7.1. ICA-AROMA

For pipelines A5 through A8, ICA-AROMA was applied immediately following the first-stage preprocessing steps described above. Given that this is a relatively high motion sample, we used aggressive denoising but otherwise default options. The ICA-AROMA code was modified to warp to MNI space via our ANTs transformation matrix rather than a FNIRT transformation.

#### 2.7.2. Temporal filtering

Following the common preprocessing steps (pipelines A1 through A4) or ICA-AROMA (A5 through A8), all pipelines underwent bandpass temporal filtering (0.01 – 0.08 Hz) done via a fast Fourier transformation. In follow up comparisons we tested the effect of a highpass filter; see *section 2.9.1. Filtering*. To avoid reintroducing artifacts, HMPs were also filtered, and WM, CSF, and the global signal were calculated following temporal filtering (Lindquist et al., 2019).

#### 2.7.3. Nuisance regression

Following filtering, nuisance regression was applied. All pipelines, including ICA-AROMA pipelines, included WM + CSF regression. Pipelines A1 through A4 included regressing the six HMPs, while pipelines A2, A4, A6 and A8 included GSR. For all nuisance parameters, both linear and quadratic terms were included, along with the first temporal derivative of those terms (4 regressors per parameter). Censoring was carried out as part of the regression step for pipelines A3, A4, A7 and A8. We censored volumes above a FD threshold of 0.25 mm (based on FSL *MCFLIRT*, i.e. FD_Jenkinson_ as described in Ciric et al., 2018), and censored only the identified frames.

### 2.8. Connectome generation

Following the above preprocessing steps, each scan was registered to the study specific template using the previously generated ANTs transformation matrix. Each voxel was then assigned to one of 325 nodes (regions) within the MIST 325 parcellation (Urchs et al., 2019). The mean time course was calculated for each node by averaging the time courses for all voxels within the node. The Pearson correlation between each pair of nodes was then calculated, generating 52650 edges. Correlation values were Fisher z-transformed to better approximate a normal distribution.

### 2.9. Pipeline comparison metrics

#### 2.9.1. Motion correlated edges (QC-FC)

Typically, preprocessing benchmark studies have looked at the amount of noise remaining in a dataset following preprocessing by QC-FC, which is the correlation between the strength of a given network edge across participants, with participant motion (as measured by average framewise displacement) (Satterthwaite et al., 2013; Power et al., 2012). QC-FC is used to examine the quantity of network edges where FC significantly relates to motion (which should theoretically be zero, notwithstanding edges functionally involved in motion); better preprocessing pipelines will have fewer edges with a significant relationship with head motion. Previous work has also assessed whether a network edge’s QC-FC is related to the inter-node distance. This addresses whether short-range connections are more vulnerable to head-motion than long-range connections, and if preprocessing steps alter the slope of this relationship (Power et al., 2015).

Using the approach of Satterthwaite et al. (2013), for each edge the correlation was calculated between scan motion (calculated as the average relative framewise displacement, as estimated from the first FSL *MCFLIRT* step) and edge strength (i.e. the Fisher z-value), across all 112 scans. Due to the concern that preprocessing steps such as GSR affects edges differently depending on the edge-length, we also assessed distance-dependent effects by plotting the edge strength-motion correlation vs. Euclidian edge length for each edge (Satterthwaite et al., 2013). Euclidian edge length was found by calculating the distance between the center of mass of the two nodes, as defined by the MIST parcellation.

Three QC-FC metrics were extracted: (1) the percentage of edges correlated with head motion at a p-value <0.05 uncorrected; (2) the mean of the absolute value of the correlation between head motion and edge strength; absolute value was used to assess the magnitude of the motion-effect, rather than the direction; and (3) the slope of the best-fit line of edge correlation with motion vs Euclidian edge length.

#### 2.9.2. Fingerprinting

While measures such as QC-FC can estimate noise removed during preprocessing, there is a concern that pipelines that aggressively remove noise may also be removing signal of interest (Ciric et al., 2017). To assess this, we used functional connectome fingerprinting to gauge whether information unique to an individual remains after preprocessing. Following the approach used by Finn et al. (2015) and related work (Byrge and Kennedy, 2019; Kaufmann et al., 2017; Miranda-Domingeuz et al., 2018), connectomes were correlated for each pair of scans. In this context, a scan’s “complement” is the other scan from the same individual. For example, the complement to the scan for child 1’s initial scan is child 1’s 12-month follow up scan, and vice versa. A scan from child 2 would be a “non-complement” scan.

The fingerprinting match rate is calculated by comparing each scan against every other scan and dividing the number of times a scan’s highest correlation is its complement by the total number of scans. Here we had 56 individuals, each with two scans, for a denominator of 112. We expressed this as a percentage. For each scan we also calculated the fingerprinting margin by taking the difference in correlation between a scan and its complement, and the highest correlation to a non-complement. This metric is indicative of the degree of individualization. We also calculated the mean correlation between complement scans and mean correlation between all non-complement scans.

To assess associations between individualization and head motion across pipelines, we plotted the fingerprinting margin vs. head motion, then calculated the slope, intercept, and R^2^. For each scan-complement pair, we used the higher of the two motion values for this analysis.

#### 2.9.3. Intersubject correlation

Because data was collected while participants watched the same set of videos, our data offered the possibility to simultaneously assess the impact of preprocessing on functional connectivity and task-evoked responses. To this end, we correlated the time series for each node between all pairs of scans to find the intersubject correlation (ISC; Hasson et al., 2004). Preprocessing pipelines that fail to adequately remove noise, or remove task-evoked signal will have lower ISC values. For pipelines that included censoring, we did not include time points that were censored for either scan in a given pair. From there, we calculated the mean ISC for each node within each scan as the average of that individual’s correlations to all other individuals. Since ISCs have significant correlations in particular brain regions (Hasson et al., 2004; Kauppi et al., 2010), such as the visual and auditory networks, we identified the 10 nodes with the highest average ISCs across all pipelines and used the average of these 10 nodes to generate a single ISC score for each scan for each pipeline. While many ISC studies only use one scan per participant (Vanderwal et al., 2019), our analysis considers how the similarity between scans might be impacted by preprocessing decisions, and we therefore included both scans per child.

#### 2.9.4. Intra-scan inter-pipeline correlation

Preprocessing choices may have relatively large or relatively small effects on connectomes, and these may influence downstream analyses and convergence across studies. To assess the magnitude of effects of preprocessing choices on FC estimates, for each scan we calculated the correlation between edge strengths across pipelines A1-A8. These were Fisher z-transformed, averaged across scans, then converted back to correlations for ease of comparison.

### 2.10. Follow-up comparisons

#### 2.10.1. Filtering

Using a lowpass filter has become common in the preprocessing literature due to frequencies above approximately 0.1 Hz being more highly associated with noise than signal of interest (Satterthwaite et al., 2019). However, this remains controversial, as connectivity information at higher frequencies will be lost (Niazy et al., 2011). We therefore repeated our analysis using a highpass (>0.01 Hz) rather than a bandpass filter, allowing us to compare the same eight pipelines. All other preprocessing steps were identical.

#### 2.10.2. Varying censoring parameters

Following our initial comparison of pipelines, we conducted an additional comparison focusing specifically on the effect of censoring at different thresholds. There is variation in thresholds used in the literature, with older studies generally using more lenient censoring than more recent studies (Satterthwaite et al., 2019). Similarly, the literature has advocated both censoring a single volume per motion artifact (Satterthwaite et al., 2013) and censoring multiple volumes per motion artifact (Power et al., 2012). Thresholding decisions are challenging because there is limited guidance in pediatric populations. Stricter censoring thresholds will reduce the number of time points, which in turn can result in less reliable functional connectivity estimates (Gordon et al., 2017).

We used pipeline A4 above, which performed well on all metrics, and modified the approach to censoring using 11 additional pipelines that varied in the censoring threshold and the volumes censored per motion artifact (Table 2). Pipelines that censor 3 volumes per motion artifact censor volumes immediately before and after any region of motion, despite neither reaching the FD threshold for censoring. When two or more volumes in a row both meet the FD threshold for censoring, only 2 additional volumes are censored, one before and one after the motion region. Similarly, for pipelines that censor 4 volumes per motion artifact, 1 volume prior and 2 volumes after the motion-contaminated volume(s) are censored. The same 56 participants (112 scans) were included across comparisons, even if stricter censoring caused scans to fall below the threshold of 11 minutes of uncensored data.

**Table 2:** Preprocessing pipelines tested to compare effects of censoring at different thresholds and volumes censored per motion artifact

| Pipeline | Volumes censored per artifact | Censoring threshold (mm FD) |
| --- | --- | --- |
| B1 | 1 | 0.3 |
| B2 (A4) |  | 0.25 |
| B3 |  | 0.2 |
| B4 |  | 0.15 |
| B5 | 3 | 0.3 |
| B6 |  | 0.25 |
| B7 |  | 0.2 |
| B8 |  | 0.15 |
| B9 | 4 | 0.3 |
| B10 |  | 0.25 |
| B11 |  | 0.2 |
| B12 |  | 0.15 |

## 3. Results

### 3.1. Participant characteristics

The mean age of participants for their baseline scan was 5.47 years old (standard deviation: 0.76 years), and 6.52 for their second scan. Time from initial to follow up scan ranged from 0.88 to 1.19 years (mean: = 1.05 years, SD = 0.069 years). The average motion of our sample was high, with a mean average relative FD of 0.126 mm across all 112 scans (median mean relative FD of 0.095 mm). The correlation in mean motion (average relative FD) between scans from the same participant was not significant (r = 0.18, p = 0.19). Across the 112 scans, age and mean FD were not significantly correlated (r = −0.14, p = 0.14).

For analyses that divided scans into two groups based on median head motion, the 56 low motion scans had a mean average FD of 0.062 mm (range: 0.035 to 0.094 mm, std = 0.016 mm). The 56 high motion scans had a mean average FD of 0.191 mm (range: 0.097 to 0.506 mm, std = 0.088 mm). Mean age at time of scan was similar in both groups (low motion: 6.01 years, std = 0.920 years; high motion: 5.98 years, std = 0.935 years).

### 3.2. QC-FC

QC-FC associations were assessed for each pipeline (Figure 1). Figure 1a shows the percentage of edges significantly correlated with motion, ranging from 42% to 20%. Both GSR and censoring reduced the number of edges correlated with motion, and pipelines that paired these two steps had the smallest number of motion-associated edges. Pipelines A1 (regress HMP) and A5 (ICA-AROMA), lacking both GSR and censoring, had 42% and 40% of edges correlated with motion respectively, which dropped to 20% (A4) and 21% (A8) respectively with the inclusion of combined GSR and censoring. In general, ICA-AROMA pipelines fared better than comparable pipelines that regressed HMP, though the pipeline with the fewest QC-FC associated edges (A4 – regress HMP+GSR+censoring) did not use ICA-AROMA.

**Figure 1.**
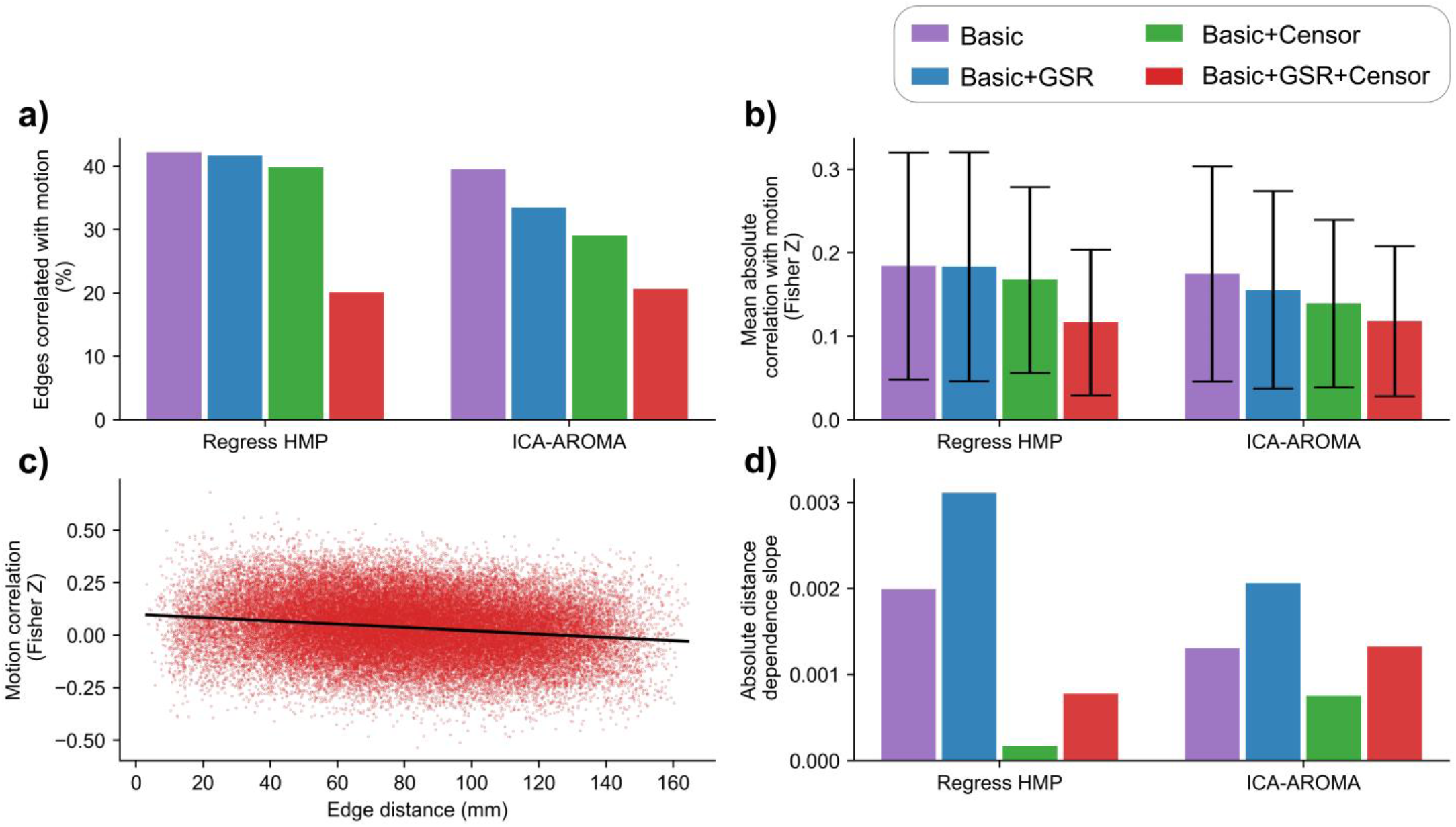
Quality control-functional connectivity (QC-FC) across pipelines. Plots are arranged from pipeline A1 through A8, as described in Table 1. **a)** Percentage of edges with a significant correlation between edge strength and subject motion across all 112 scans (uncorrected p < 0.05). **b)** Mean and standard deviation of absolute correlation between edge strength and subject motion across all 112 scans. **c)** Example QC-FC distance dependence plot, from pipeline A4 (regress HMP + GSR + censor). Each point is an edge in the connectome, plotted based on the length between its nodes (edge distance) and correlation between edge strength and subjection motion. **d)** Absolute value of the slope of each pipeline’s distance-dependence plot.

Figure 1b shows the mean absolute correlation with motion for all edges. Here the same trends are observed, with GSR and censoring reducing the effect of motion. A1 and A5 had mean absolute correlations with motion of z = 0.184 and z = 0.175 respectively. When adding both GSR and censoring (A4 and A8), these dropped to z = 0.117 and z = 0.118 respectively, with only smaller improvements when using only one of GSR or censoring.

Figure 1c shows an example from pipeline A4 of edge correlation with motion vs. edge distance with the slope indicated. Figure 1d shows the slope values for each pipeline. Our results agreed with previous literature that GSR is associated with a more negative slope, suggesting GSR affects shorter edges differently than longer edges, with shorter edges having more remaining motion influence, but longer edges more likely to be negatively correlated with motion. For example, A1 (regress HMP+no GSR+no censoring) had a slope of −0.0020, which increased to −0.0031 when adding GSR (A2). However, we found that censoring reduces the slope, partially compensating for the effect of GSR. The two pipelines with the smallest slope were A3 (regress HMP+no GSR+censoring), with a slope of −0.0002 and A4 (regress HMP+GSR+censoring) with a slope of −0.0008.

When scans were split into lower- and higher-motion subsets, the higher-motion scans fared worse on all metrics (Figure 2a vs 2b, 2c vs 2d, 2e vs 2f). Both the lower- and higher-motion groups had fewer edges significantly correlated with motion than the entire group (Figures 2a and 2b vs 1a), which we attribute to a smaller sample size and a smaller range of possible motion values. In the low motion group, the choice of pipeline had minimal effect on the percent of edges correlated with motion (ranging from 6.1% to 8.9%; Figure 2a) and the mean absolute correlation with motion (ranging from z = 0.11 to z = 0.13; Figure 2c). Small improvements were seen with censoring, and pipelines that included GSR fared slightly worse. Most of the effects noted in the entire sample (Figure 1) were only seen in the high motion group (Figures 2b and 2d), where censoring and GSR used together improve the percentage of edges significantly correlated with motion (e.g. pipeline A1 to A4 dropping from 20% to 11%) and the mean absolute correlation with motion (e.g. A1 to A4 dropping from z = 0.17 to 0.13). Interpipeline slope trends were similar for both lower- and higher-motion groups as in the entire sample (2e and 2f vs 1d), with GSR making the slope more negative, but censoring mitigating the effect.

**Figure 2.**
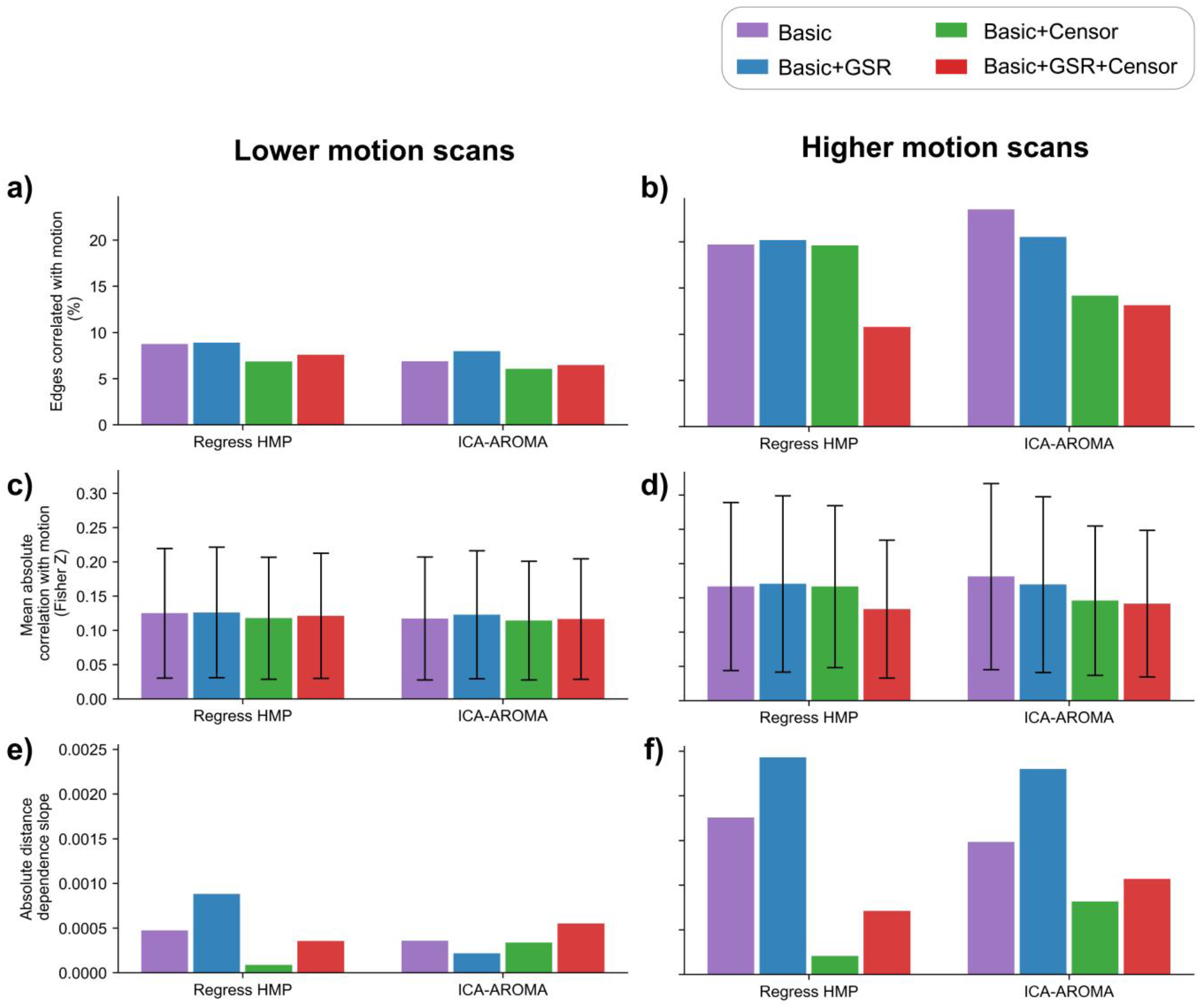
Quality control-functional connectivity (QC-FC) across pipelines, separately for lower or higher motion scans. Higher and lower motion participants were identified as above or below median average framewise displacement. Plots are arranged from pipeline A1 through A8, as described in Table 1. **a, b)** Percentage of edges that have a significant correlation between edge strength and subject motion (uncorrected p < 0.05). **c, d)** Mean and standard deviation of absolute correlation between edge strength and subject motion. **e, f)** Absolute value of the slope of each pipeline’s distance-dependence plot.

### 3.3. Fingerprinting

Fingerprinting match-rate was assessed for each pipeline (Figure 3a). Overall, the match rate was high for all pipelines tested, ranging from 71% to 90% (chance level is <1%). GSR had a minimal effect on the overall match rate (e.g. 71.4% for A1 – regress HMP+no GSR+no censor, and 70.5% for A2 – regress HMP+GSR+no censor), but pipelines that included censoring were more successful. The two pipelines with the highest match rate included censoring and regressing HMP rather than using ICA-AROMA (A3 – no GSR and A4 - GSR; 89.3% and 90.2% respectively).

**Figure 3.**
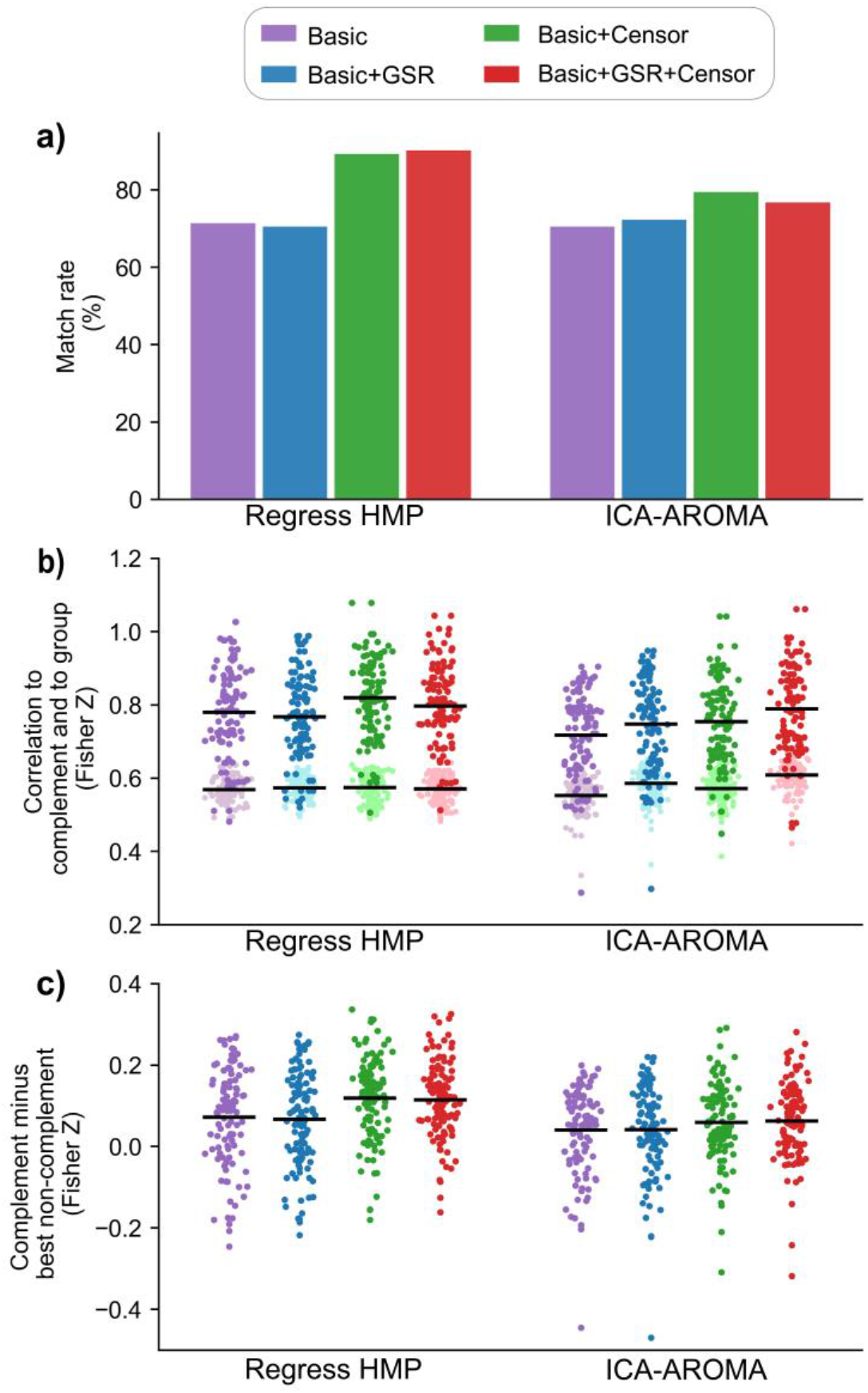
Functional connectome fingerprinting by pipeline. Plots are arranged from pipeline A1 through A8, as described in Table 1. **a)** Match rate across pipelines. A scan matched if its highest correlation was to its complement scan (other scan from the same individual). **b)** Each scan’s correlation to their complement scan (darker points) and their average correlation to every other scan (lighter points), for each pipeline. Lines represent mean values. **c)** Fingerprinting margin, i.e. difference between each scan’s correlation to their complement and to their best (highest correlation) non-complement. Any point below 0 fails to successfully match. Lines represent mean values.

Scans’ mean correlation to non-complement scans varied minimally between participants and across pipelines (Figure 3b), suggesting group network structure can be consistently found regardless of preprocessing choices. While the correlation between a scan and its complement showed a much larger range of values, this is likely an artifact of being a single correlation, rather than an average. The two pipelines with the highest fingerprinting match-rate, A3 and A4, were the ones with the highest mean correlation between scan-complement pairs (A3 – z = 0.82 and A4 – z = 0.80), though interestingly pipeline A8 (ICA-AROMA+GSR+censoring) had a comparable mean correlation between scan-complement pair (z = 0.79), though a far larger mean correlation to non-complement scans (z = 0.61, vs z = 0.57 for both A3 and A4).

Figure 3c shows the fingerprinting margin (i.e. complement minus best non-complement) for each of the 112 scans. As expected, pipelines with a higher match-rate (Figure 3a) have a higher average fingerprinting margin, which ranged from z = 0.040 to z = 0.072 for the less successful pipelines (A1, A2, A5 through A8), while reaching z = 0.12 and z = 0.11 for A3 and A4 respectively. The group distribution for this metric was relatively consistent, with more successful pipelines shifting the distribution upwards. This suggests that with certain pipelines scans that fail to match with their complement still see greater recovery of individual information via a reduced margin of failure (e.g. a scan’s fingerprinting margin may improve from −0.2 to −0.1), and scans that successfully match pass with an improved margin (e.g. improving from +0.1 to +0.2).

Figure 4 shows the fingerprinting margins (from Figure 3c) plotted against head motion. A more negative slope suggests that the pipeline performs progressively worse at identifying individuals with more motion, and a higher r^2^ value suggests that data run through the pipeline is more sensitive to motion noise. In general, pipelines that regress HMP rather than use ICA-AROMA have less negative slopes, lower r^2^ values, and higher intercepts. For example, pipeline A1 (regress HMP+no GSR+no censor) has a slope = −0.51, intercept = 0.20, r^2^ = 0.36, while A5 (ICA-AROMA+no GSR+no censor) has slope = −0.62, intercept = 0.19, r^2^ = 0.41. Pipelines that include GSR have more negative slopes and higher r^2^ values, for example going from A1 to A2 increases the slope from −0.51 to −0.56, and r^2^ increases from 0.36 to 0.40. Censoring has minimal effect on slope, but raises the intercept and lowers the r^2^, for example going from A1 to A3 raises the intercept from 0.20 to 0.22, and lowers r^2^ from 0.36 to 0.25. These suggest that censoring benefits all scans, but GSR has minimal benefit for fingerprinting, especially at higher motion. The participant with the most motion (mean relative FD of 0.51 mm for one of their two scans) could be considered an outlier – repeating the analysis with that participant removed reduces the r^2^ and makes the slopes less negative, but trends between pipelines remain (Supplementary Figure 1).

**Figure 4.**
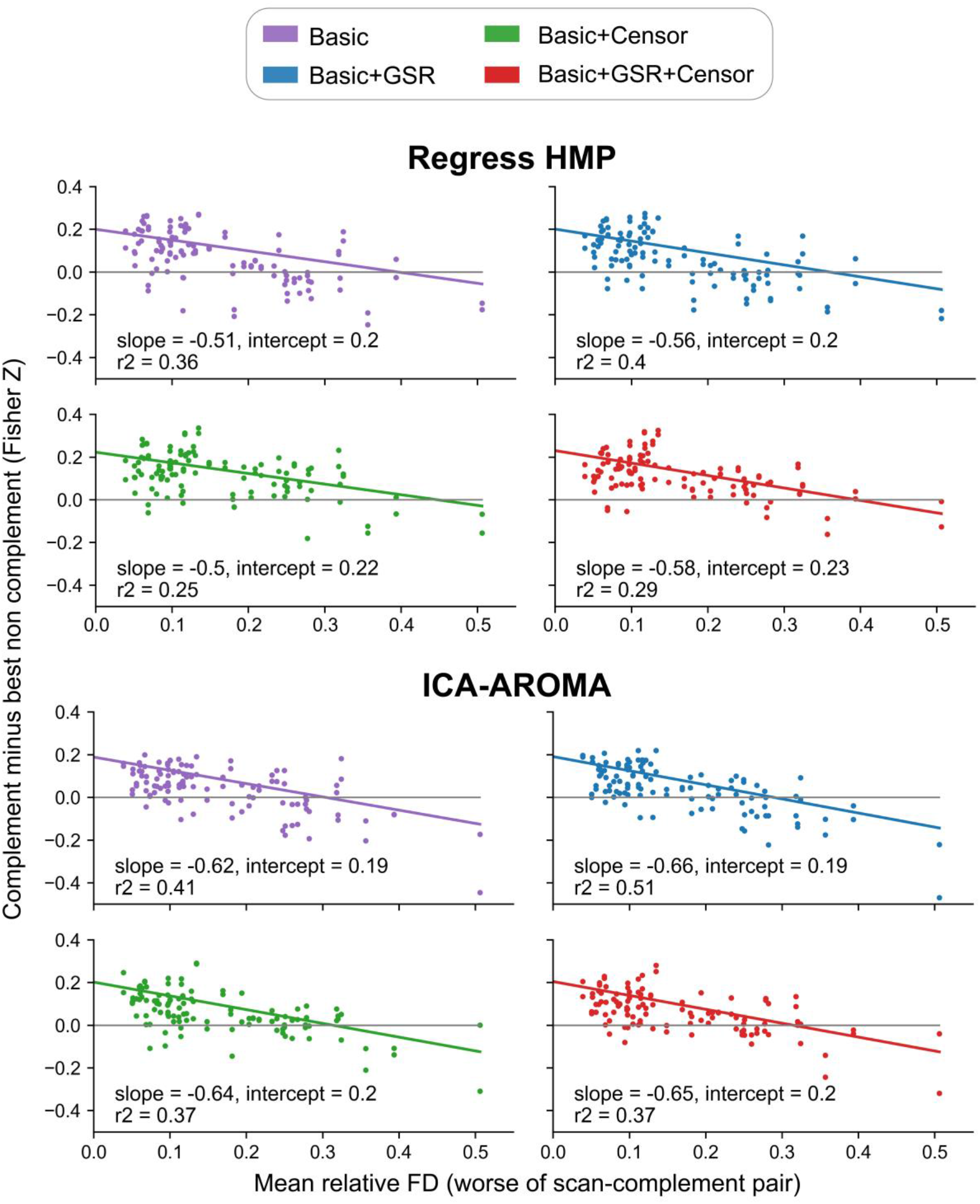
Fingerprinting margin as a function of head motion. The difference between each scan’s correlation to their complement scan minus their correlation to the best (highest correlation) non-complement, plotted by motion. Motion was calculated by taking the worse of each participant’s two scans’ mean relative framewise displacement; each participant has two points for their two scans. Any point below 0 on the y axis fails to successfully match. Plots are arranged, left to right, from pipeline A1 through A8, as described in Table 1.

### 3.4. Intersubject correlation

Figure 5 shows each participant’s mean intersubject correlation scores (ISCs) across pipelines, based on the 10 parcels with the highest scores. Both GSR and censoring improved ISCs, incrementally when applied together. Pipelines A1 (regress HMP) and A5 (ICA-AROMA), both lacking GSR and censoring, had mean ISCs of z = 0.16 and z = 0.17 respectively, which increased to z = 0.21 for both pipelines when both strategies were added (pipelines A4 and A8). Pipelines that used ICA-AROMA had slightly higher ISCs than pipelines that did not, though the gain was small.

**Figure 5.**
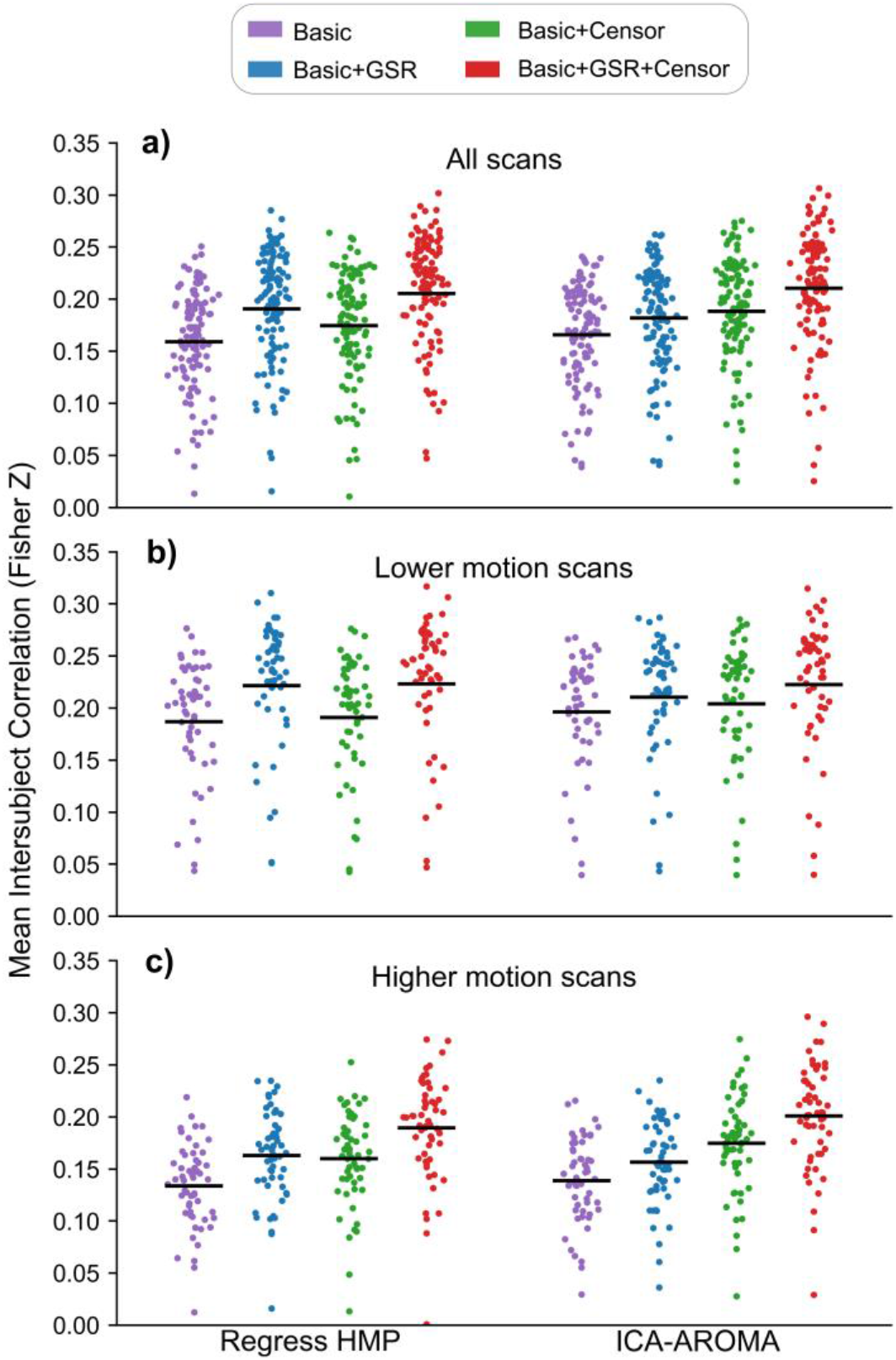
Mean intersubject correlation (ISC) values. Each point represents the mean ISC for a scan to all other scans, averaged across the 10 parcels with highest ISC. Lines represent mean values. Plots are arranged from pipeline A1 through A8, as described in Table 1. **a)** Mean ISCs for all 112 scans, each compared to all other scans. **b)** Mean ISCs for each of the 56 below-median-motion scans, compared to the other 55. **c)** Mean ISCs for each of the 56 above-median-motion scans, compared to the other 55.

When only low motion scans were compared using ISC (Figure 5b) the benefits of censoring were minimal, with or without GSR. GSR itself increased ISCs, for example A2 (regress HMP+GSR+no censor) had a mean ISC of z = 0.22, compared with z = 0.19 for A1 (regress HMP+no GSR+no censor). However, when only high motion scans were compared (Figure 5c), the benefits of censoring were more apparent, especially in combination with GSR. For example, pipeline A2 (regress HMP+GSR+no censor) had a mean ISC of z = 0.16, which increased to z = 0.19 for A4 (regress HMP+GSR+censor).

### 3.5. Intrascan inter-pipeline correlation

Figure 6 shows the within-scan between-pipeline correlation of whole brain functional connectivity estimates. These correlations were converted to z-scores for averaging, then converted back to correlations for ease of interpretability. While correlations were high overall, ranging from 0.67 to 0.97, a correlation as low as 0.67 for the same scans preprocessed in two different ways (in this case between pipelines A3 (regress HMP+no GSR+censor) and A6 (ICA-AROMA+GSR+no censor) highlights concerns about replicability given the overall influence of preprocessing.

**Figure 6.**
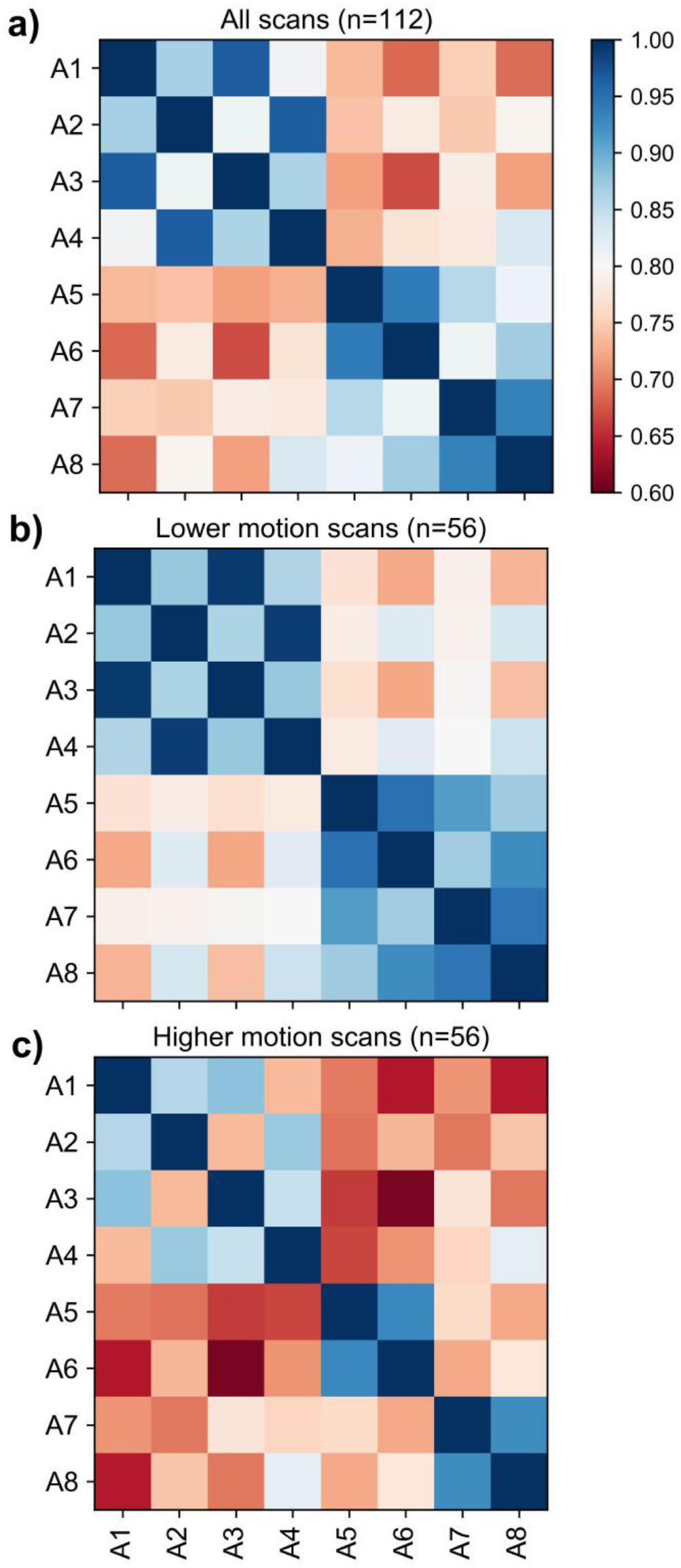
Intrascan inter-pipeline correlations. For each scan, the correlation between edge strengths was compared across pipelines, Fisher z-transformed, averaged across scans, then converted back to correlations. Pipelines are listed in Table 1, briefly: A1 – regress HMP; A2 – regress HMP + GSR; A3 – regress HMP + censor; A4 – regress HMP + GSR + censor; A5 – ICA-AROMA; A6-ICA-AROMA + GSR; A7 – ICA-AROMA + censor; A8 – ICA-AROMA + GSR + censor. **a)** Average correlation based on all 112 scans. **b)** Average correlation using the 56 scans below the median average framewise displacement. **c)** Average correlation using the 56 scans above the median average framewise displacement

Figure 6 also shows how each preprocessing choice progressively changes functional connectivity estimates. When comparing pipelines that were identical but either included GSR or not, correlations were 0.87, 0.86, 0.94 and 0.94 (for comparisons A1-A2, A3-A4, A5-A6, A7-A8 respectively). When comparing pipelines that were identical but either included censoring or not, correlations were 0.97, 0.96, 0.85, and 0.87 (for comparisons A1-A3, A2-A4, A5-A7, A6-A8 respectively). Individually these changes were small but comparing pipelines A1 and A5 to A4 and A8, respectively, i.e. neither GSR nor censoring, and with both GSR and censoring, the correlation was only 0.81 for both comparisons. The choice to use ICA-AROMA vs. regressing HMP had a larger effect, with correlations ranging from 0.74 to 0.83. Again, these correlations decreased further if the pipelines also differed in use of GSR or censoring. Unsurprisingly, preprocessing choices had a smaller effect on low motion scans (Figure 6b), with correlations between pipelines ranging from 0.72 to 0.99. For high motion scans (Figure 6c), correlations ranged from 0.61 to 0.93.

### 3.6. Filtering comparison

When a highpass filter was used rather than a bandpass filter, overall trends between pipelines were quite similar (see Supplemental Figures 2–4). With a highpass filter, the number of edges significantly correlated with motion ranged from 25-50% (Supplemental Figure 2a). Other than for pipeline A5 (which fared poorly on this metric regardless), bandpass filtering resulted in fewer edges significantly correlated with motion than highpass filtering, for example 20% vs 27% of edges for pipeline A4 (regress HMP+GSR+censor). Similarly, pipelines that used bandpass filtering had a smaller mean absolute correlation with motion (Supplemental Figure 2b), for example z = 0.12 vs z = 0.14 for A4. Highpass filtering resulted in more negative edge correlation with motion vs. edge distance slopes than bandpass filtering (e.g. −0.0017 vs −0.0008 for A4; Supplemental Figure 2c), suggesting a bandpass filter leads to reduced distance dependent effects.

For fingerprinting, running the analysis with a highpass filter usually led to a lower fingerprinting match rate (Supplemental Figure 3a), such as 65% vs 71% for pipeline A2 (regress HMP+GSR+no censor). However, pipelines A3 (regress HMP+no GSR+censor) and A4 (regress HMP+GSR+censor) were equally successful using either style of filter (89% match rate for A3, 90% for A4). Pipelines that used highpass filtering showed increased correlations to both non-complement scans and to complement scans (Supplemental Figure 3b) (e.g. for A4: mean complement correlation z = 0.87 for highpass, z = 0.80 for bandpass), suggesting a highpass filter makes all scans’ FC estimates more similar to each other, relative to bandpass. This suggests that signals above 0.08 Hz, whether noise or connectivity information, are more consistent across people and longitudinally within people. Filtering choice had a variable effect on the fingerprinting margin (Supplemental Figure 3c). For some pipelines, using a highpass filter led to a lower mean complement minus non-complement score (e.g. A2, z = 0.044 for highpass, z = 0.067 for bandpass), but for pipelines A3 and A4 this margin of fingerprinting success improved with a highpass filter (e.g. A4, z = 0.12 for highpass, z = 0.11 for bandpass).

Bandpass filtering increased ISCs for all pipelines compared to highpass filtering (Supplemental Figure 4) (e.g. A4, z = 0.18 for highpass, z = 0.21 for bandpass), suggesting that bandpass filtering is effective for removing noise while retaining task signal. However, at least part of this improvement with bandpass filtering may be explained by signals composed of a narrower range of frequencies being inherently more similar.

### 3.7. Censoring comparison

We repeated our initial analysis by varying the amount of censoring with a pipeline that used a bandpass filter, regressed HMP and included GSR (Figure 6). When one volume was censored per motion artifact, a lower threshold for censoring resulted in fewer edges being correlated with motion, lowering from 21% of edges at a threshold of 0.30 mm, to 17% of edges at a threshold of 0.15 mm (Figure 6a). However, the censoring threshold had minimal effect on the number of edges significantly correlated with motion when 3 or 4 volumes were censored per artifact, ranging from 13% to 15%. These effects were repeated in the mean absolute correlation with motion (Figure 6b), with a lower threshold at one volume per artifact reducing the correlation from z = 0.119 at 0.30 mm to z = 0.109 at 0.15 mm, but having minimal effect at 3 or 4 volumes (ranging from z = 0.099 mm to 0.105 mm).

The fingerprinting match-rate decreased as censoring became more stringent (Figure 6c), suggesting that although censoring can help with identifiability, information about individuals is lost with more stringent approaches. At 1 volume censored per motion artifact, the decrease was from 91% to 88%, but at 4 volumes censored per artifact the match-rate fell from 88% to 79% when the threshold changed from 0.30 mm to 0.15 mm. With stricter censoring, both the mean correlation to a complement scan and the mean correlation to a non-complement scan decreased (Figure 6d), for example at 3 volumes per motion artifact the mean complement-correlation dropped from z = 0.784 to z = 0.731 for a threshold of 0.30 mm and 0.15 mm respectively. Surprisingly, the fingerprinting margin was largely unaffected by censoring (Figure 6e), ranging from a mean of z = 0.102 to z = 0.116, except for the most stringent censoring of 3 volumes at 0.15 mm and 4 volumes at 0.15 mm having a margin mean of z = 0.091 and z = 0.077 respectively.

**Figure 7.**
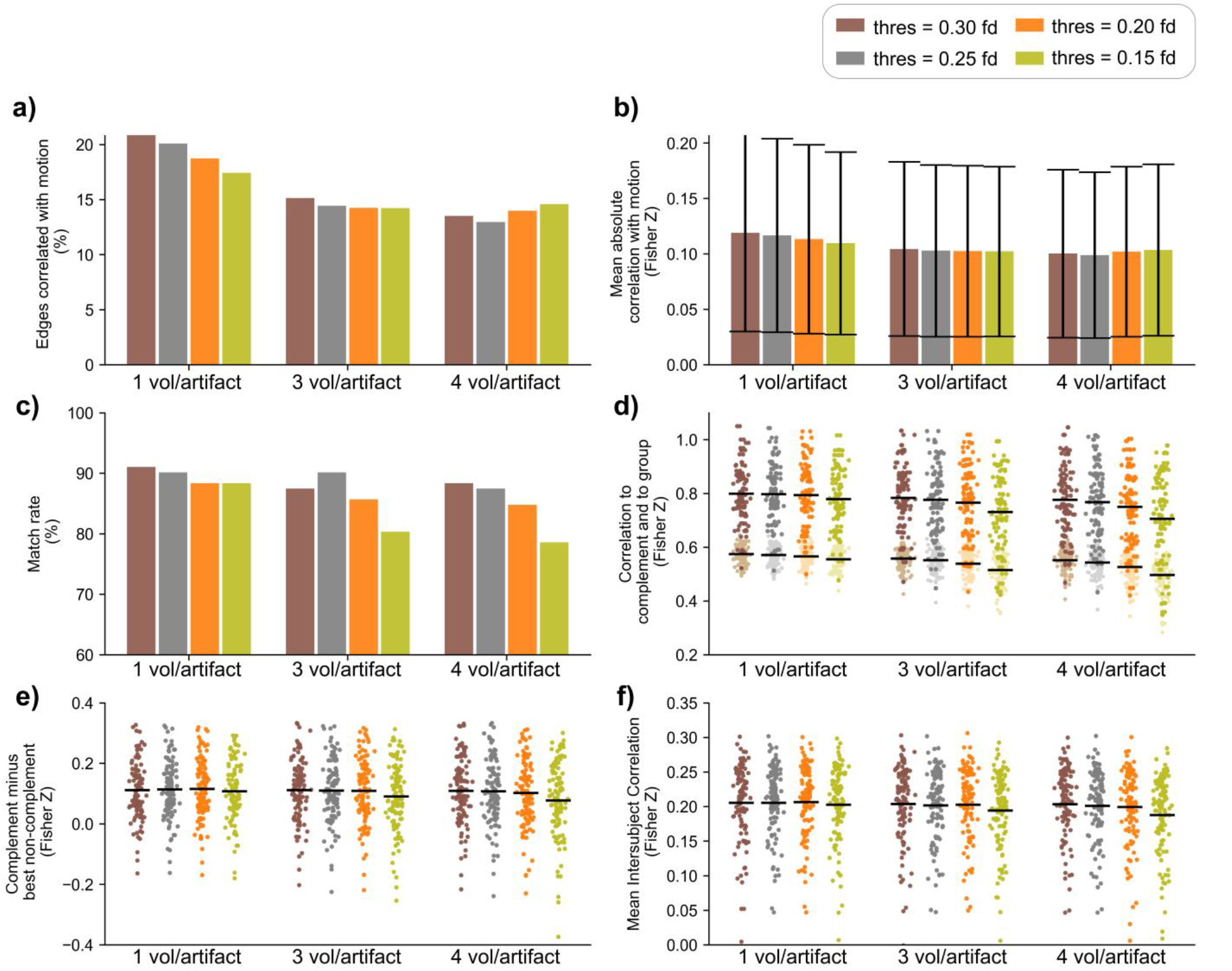
Pipeline benchmarks with different censoring thresholds. Colored bars depict four threshold levels and across the x-axis a different number of volumes is censored around the high motion frame. Plots are arranged from pipeline B1 through B12, as described in Table 2. **a)** Percentage of edges with a significant correlation between edge strength and subject motion across all 112 scans (uncorrected p < 0.05). **b)** Mean absolute correlation between edge strength and subject motion across all 112 scans. **c)** Fingerprinting match rate across pipelines. **d)** Each scan’s correlation to their complement scan (darker points) and their average correlation to every other scan (lighter points), for each pipeline. Lines represent mean values. **e)** Difference between each scan’s correlation to their complement and to their best (highest correlation) non-complement. Lines represent mean values. **f)** Mean ISCs for all 112 scans.

Censoring has a minimal effect on ISC (Figure 6f), except for the strictest censoring. At a threshold of 0.15 mm, 3 volumes per motion artifact had a mean ISC of 0.194; 4 volumes per motion artifact had a mean ISC of 0.188. All other censoring pipelines tested ranged from 0.199 to 0.207. It should be noted that for the purposes of ISC, censoring one scan necessarily censors the scan it is being compared to, since the same time points need to be compared. A slightly stricter censoring will lead to a much larger decrease in the number of shared time points, so benefits in the removal of noise may be cancelled by the decrease in data being compared.

## 4. Discussion

Here we compared several preprocessing pipelines in relatively high motion longitudinal data from young children. We found that preprocessing strategies that include GSR and censoring together outperformed other pipelines on QC-FC metrics. We used connectome fingerprinting to estimate how individually unique information was retained, or enhanced, by preprocessing steps and found that volume censoring conferred more benefit than GSR, and that the best pipelines regressed out HMPs rather than using ICA-AROMA. When examining ISC, GSR offered the most improvement, with censoring showing a specific benefit for high motion scans. Censoring multiple volumes per motion artifact improved QC-FC metrics but had minimal or even a negative effect on other metrics. When examining overall effect of preprocessing choices on connectomes, we found that pipelines had a major effect on individual estimates of FC, which is a concern for cross-study replicability.

When comparing ICA-AROMA pipelines to pipelines that regressed HMPs, we found ICA-AROMA to be less effective on most metrics. This is somewhat surprising, given that ICA-AROMA primarily works by removing independent components strongly associated with motion estimates (Pruim et al., 2015). Given that ICA-AROMA does reasonably well on QC-FC metrics, which estimate noise-contaminated edges, but relatively poorly on fingerprinting metrics, which estimate individual information, one interpretation is that ICA-AROMA is removing both noise and signal of interest. However, we note that ICA-AROMA pipelines showed no obvious difference on ISC metrics. Other implementations of ICA-based preprocessing exist and are worth systematically investigating. ICA-AROMA is often used in conjunction with ICA-FIX, another ICA-based denoising strategy (Salimi-Khorshidi et al., 2014), and ICA-AROMA can also be run with less aggressive denoising. Some studies have combined ICA-based denoising with regressing HMPs (Kaufmann et al., 2017; Jalbrzikowski et al., 2020), which may have advantages, though effectively removes aspects of the signal related to motion twice, contrary to the original intent of ICA-AROMA (Pruim et al., 2015).

While GSR remains contentious in the field, our results suggest there are advantages in a high-motion dataset such as the early childhood sample used here. GSR reduced the number of edges significantly correlated with motion, especially when applied with censoring. Similarly, while there are distance-dependent effects when GSR is used, this effect was largely mitigated in pipelines that also included censoring. GSR had no obvious impact on connectome fingerprinting, both in terms of the match rate and the fingerprinting margin. This suggests that while GSR effectively centers, and has distance-dependent effects, these changes are less edge-specific impact than censoring. The finding that GSR had a minimal impact on fingerprinting, our metric of individual information, may alleviate concerns about its use when comparing healthy controls and clinical populations, or in developmental research, even in light of research that suggests the global signal is significantly related to life outcomes and psychological function (Li et al., 2019). We also found benefits to the use of GSR in improving signal-to-noise in ISC metrics, suggesting that the benefits in terms of motion and physiological noise removal may outweigh the cost of removing some signal of interest (Behzadi et al., 2007).

We found relatively small effects of a bandpass filter compared to a highpass filter. Pipelines that used a bandpass filter fared better on both QC-FC metrics and ISC metrics. In connectome fingerprinting, while complement correlations were higher in pipelines that used a highpass filter, so too were non-complement correlations by comparable levels. This suggests that any resulting changes to the functional connectome by including higher frequencies are consistent across all scans and are unlikely to reflect individual-specific information.

We found benefits to volume censoring across metrics, but especially in connectome fingerprinting, where censoring increased accuracy more than any of the other preprocessing steps compared here. Censoring also conferred additional benefits to pipelines that used ICA-AROMA, despite ICA-AROMA being intended partially as an alternative to censoring (Pruim et al., 2015). Censoring had only a small effect on low-motion scans for ISC comparisons, but censoring increased ISC scores on high-motion scans. When only one volume was censored per motion artifact, the more stringent the censoring threshold, the fewer edges were significantly correlated with motion. However, censoring multiple volumes per motion artifact outperformed single volume censoring, suggesting that censoring more volumes per motion artifact at a less conservative threshold may be preferable. Stricter censoring tended to decrease correlations to both complement scans and non-complement scans. This suggests that removing data points removes information that is common across participants. Interestingly, the distribution of fingerprinting margin values was relatively unaffected by the censoring threshold, except when very strict censoring was used, even though the overall match rate decreased when censoring became stricter. This suggests not all scans are affected in the same way by changes in censoring parameters.

Our findings on the intrascan interpipeline correlations are concerning towards the goal of having replicable results in FC-MRI studies. While preprocessing choices have a smaller impact on low motion data, our work suggests that when analyzing high motion data (which data from early childhood tends to be), each preprocessing choice has a drastic effect on FC estimates. While more broad methodological differences or group-level participant differences are often cited as reasons for different findings between studies, we suggest that differences in preprocessing also play a role. Preprocessing choices should be both closely considered and accurately reported. While we compared the correlation between the entire functional connectome, future work should explore regional specific effects in more detail. This can both provide more insight into how preprocessing affects FC estimates, but also provide greater clarification when different studies come to different conclusions.

There are several limitations to our work. Our study was done in young children who were participating in a passive viewing task, and our results may not be directly applicable to other populations or other study protocols. Likewise, the benefits of any step we tested may not be transferable to a more traditional task-based fMRI study, especially censoring data points during the task. We note that several of the steps implemented here could have been applied in alternative forms, which also restricts generalizability. For example, we used a 24 HMP model rather than 6 or 12 HMP (Friston et al., 1996). Similarly, there are many other thresholds that can be used for censoring or temporal filtering. There are also other possible approaches to noise mitigation, such as CompCor (Behzadi et al., 2007) or ANATICOR (Jo et al., 2010), that warrant further investigation.

We also acknowledge that choices such as parcellation may have impacted our findings. Previous studies using functional connectome fingerprinting have used a similar number or fewer nodes; for example, Finn et al (2015) used a 268-node atlas while Miranda-Dominguez et al. (2018) used a 333-node atlas. Finn et al. (2015) also found lower fingerprinting success using a 68-node atlas; it is unknown if a higher number of nodes may offset the difference between preprocessing strategies on fingerprinting metrics. While we chose metrics that we believe are meaningful to gauging the effect of preprocessing, other metrics could have instead been chosen. For example, we did not consider the effect preprocessing has on the magnitude of FC estimates within a scan, and we also did not consider the effect preprocessing might have on individual networks within the broader connectome.

Here we implemented ISC by comparing parcels across scans, rather than a voxel-based approach (Hasson et al., 2004). This has the potential to average out the effect of the task, as not all voxels within a parcel will respond to the passive viewing task in a similar way. We also chose to focus on the parcels with the highest average ISC values. These choices limit the extent our findings can be generalized across the whole brain or to voxel-specific changes. Our analysis has assumed that a higher ISC value reflects greater recovery of true signal, based on the idea that unremoved noise in one or both scans will lower the temporal correlation between scans. Parkes et al. (2018) suggest that FC test-retest reliability can increase in a high noise dataset, suggesting that noise has a highly reproducible component within an individual. It is unknown whether noise could have a similar effect between subjects in the context of ISC. Future work should investigate more specifically the effect of preprocessing choices on ISC values across the brain, especially in high noise scans.

## 5. Conclusions

Our findings have implications for future work in the field, both in pediatric studies but also fMRI studies in general. While in different datasets different preprocessing choices may be more appropriate, our results suggest that GSR, censoring, a bandpass filter, and HMP regression are preferable in high motion datasets from pediatric populations engaging in a passive viewing task. In particular, GSR and censoring showed few disadvantages across our metrics, and ICA-AROMA showed no major improvement compared to regressing out HMPs, especially when used without censoring.

All preprocessing choices have unintended effects on data; in light of the major effect that preprocessing choices has on FC estimates within a single scan, we urge awareness of the effect of preprocessing choices within any given research protocol. Ideally, studies should aim to reduce head motion at the time of scan, and new preprocessing strategies or other advances in fMRI protocols should aim to improve the signal-to-noise ratio with fewer tradeoffs.

## CRediT authorship contribution statement

**Kirk Graff:** Conceptualization, Methodology, Validation, Formal analysis, Investigation, Writing - original draft, review & editing, Visualization, Funding acquisition. **Ryann Tansey:** Investigation, Writing - review & editing. **Amanda Ip:** Resources, Investigation. **Christiane Rohr:** Investigation, Writing - review & editing. **Dennis Dimond:** Investigation, Writing - review & editing. **Deborah Dewey:** Methodology, Writing - review & editing. **Signe Bray:** Conceptualization, Investigation, Methodology, Supervision, Project administration, Funding acquisition, Writing - review & editing.

## Acknowledgments

We thank all the children and their families who participated in this study, and the staff at the Alberta Children’s Hospital Imaging Centre. Thanks to Taylor Johnson for assistance with the artwork.

## Appendix Supplementary materials

**Supplemental Figure 1.**
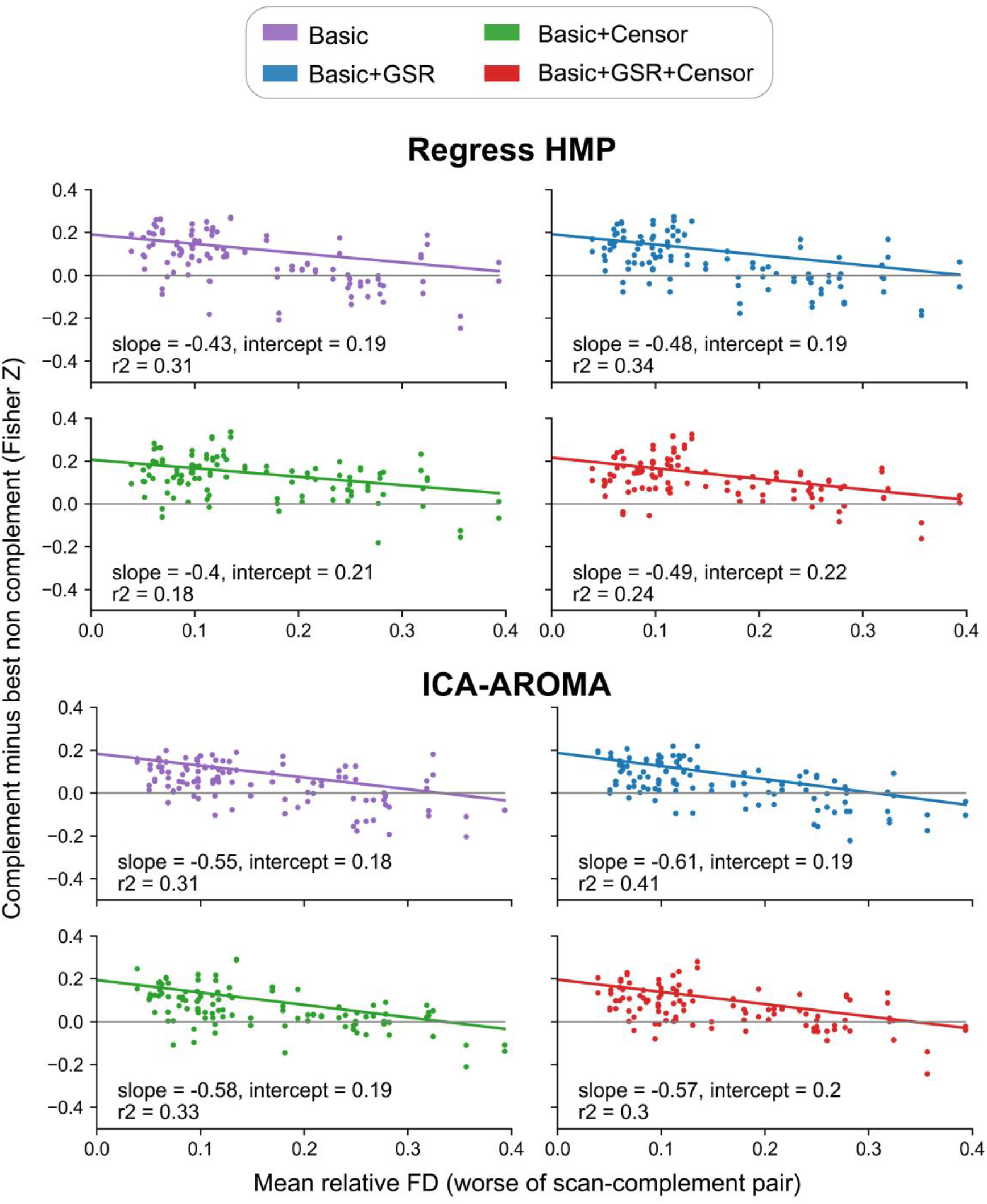
Fingerprinting margin as a function of head motion. This is similar to Figure 4 with an outlier removed. The difference between each scan’s correlation to their complement scan minus their correlation to the best (highest correlation) non-complement, plotted by motion. Motion was calculated by taking the worse of each participant’s two scans’ mean relative framewise displacement; each participant has two points for their two scans. Any point below 0 on the y axis fails to successfully match. Plots are arranged, left to right, from pipeline A1 through A8, as described in Table 1.

**Supplemental Figure 2.**
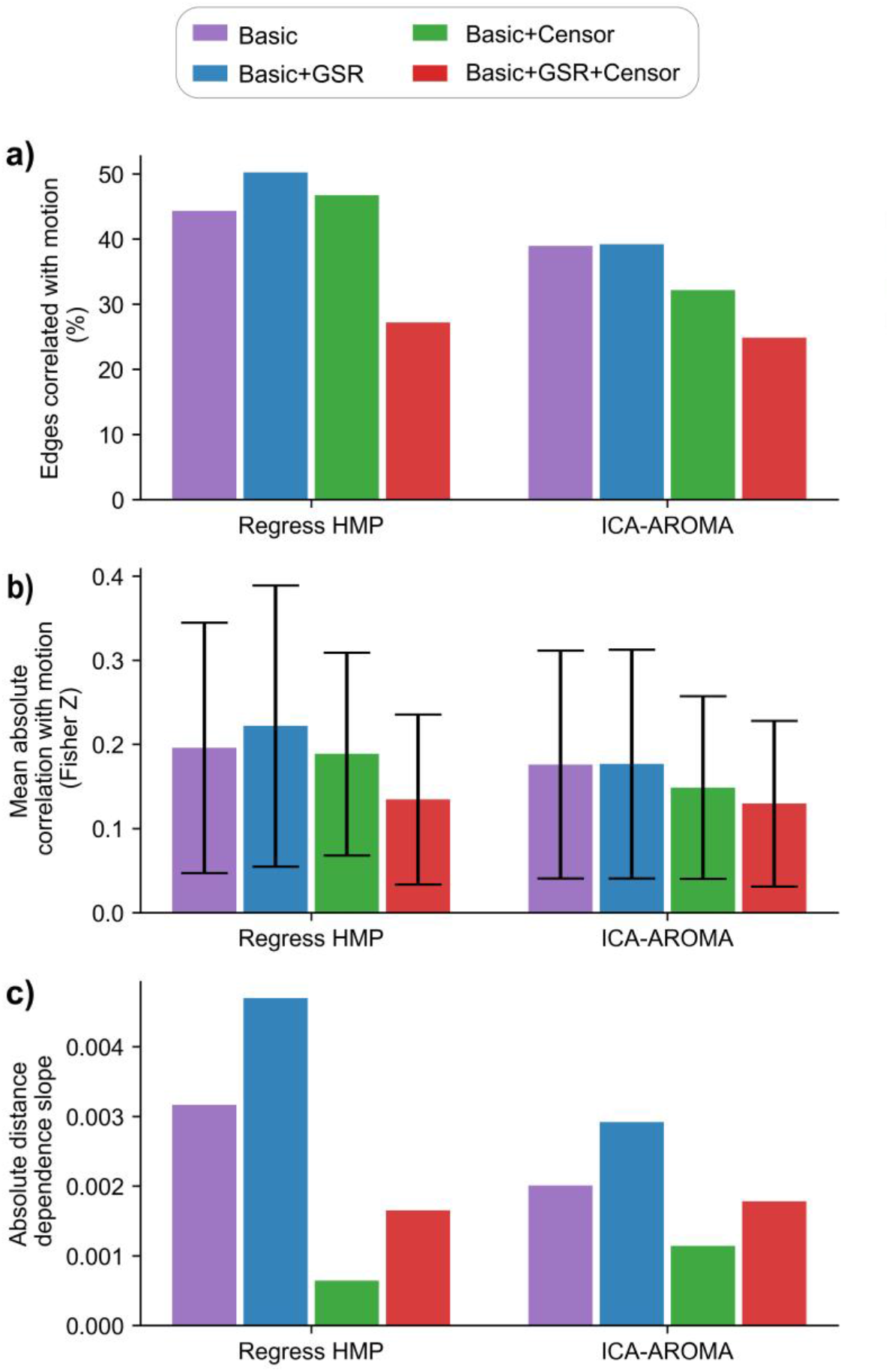
Quality control-functional connectivity (QC-FC) across pipelines with a highpass filter. Plots are arranged from pipeline A1 through A8, as described in Table 1. **a)** Percentage of edges with a significant correlation between edge strength and subject motion across all 112 scans (uncorrected p < 0.05). **b)** Mean absolute correlation between edge strength and subject motion across all 112 scans. **c)** Absolute value of slope of each pipeline’s distance-dependence plot.

**Supplemental Figure 3.**
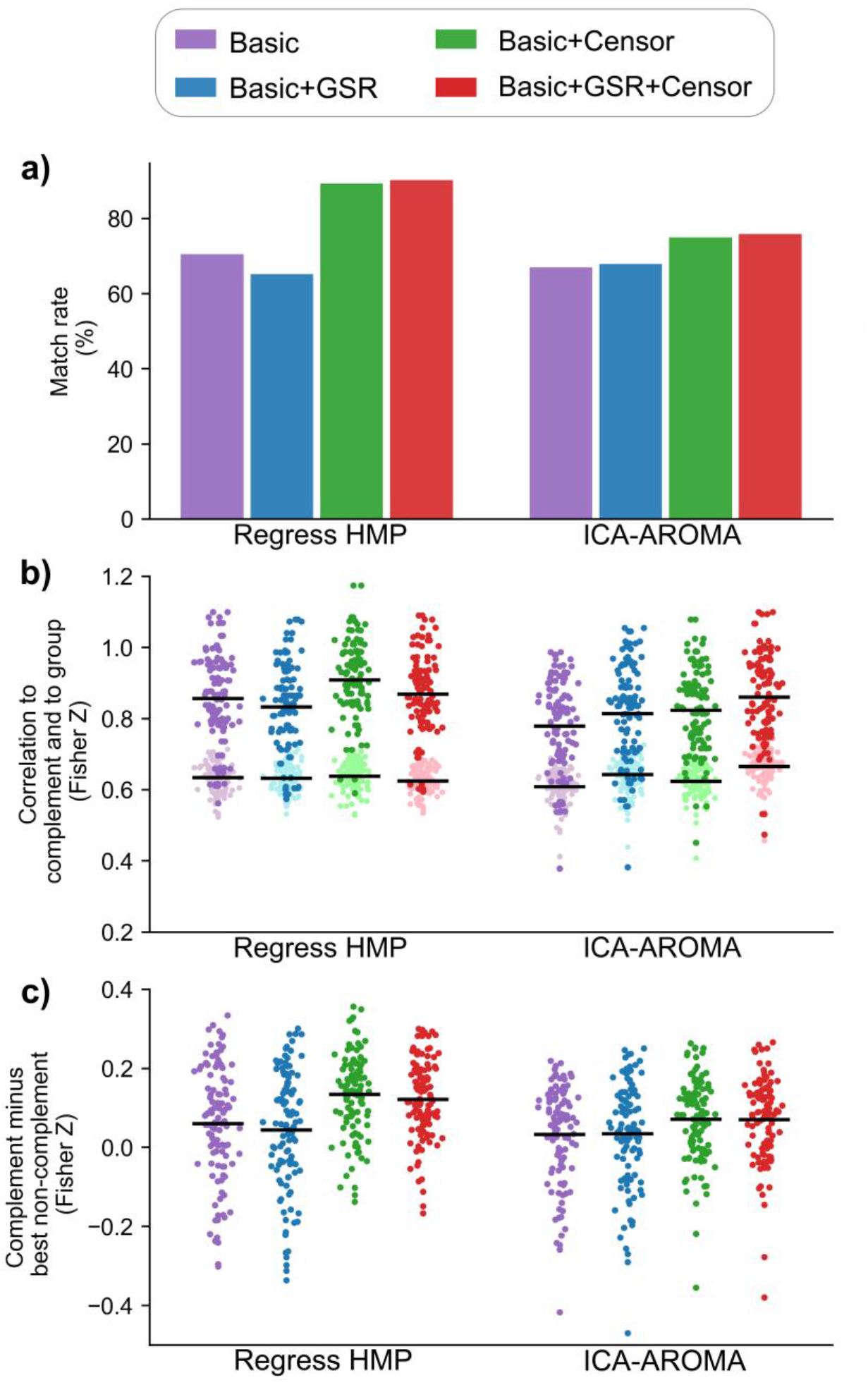
Functional connectome fingerprinting by pipeline with a highpass filter. Plots are arranged from pipeline A1 through A8, as described in Table 1. **a)** Match rate across pipelines. A scan matched if its highest correlation was to its complement scan (other scan from the same individual). **b)** Each scan’s correlation to their complement scan (darker points) and their average correlation to every other scan (lighter points), for each pipeline. Lines represent mean values. **c)** Difference between each scan’s correlation to their complement and to their best (highest correlation) non-complement. Lines represent mean values.

**Supplemental Figure 4.**
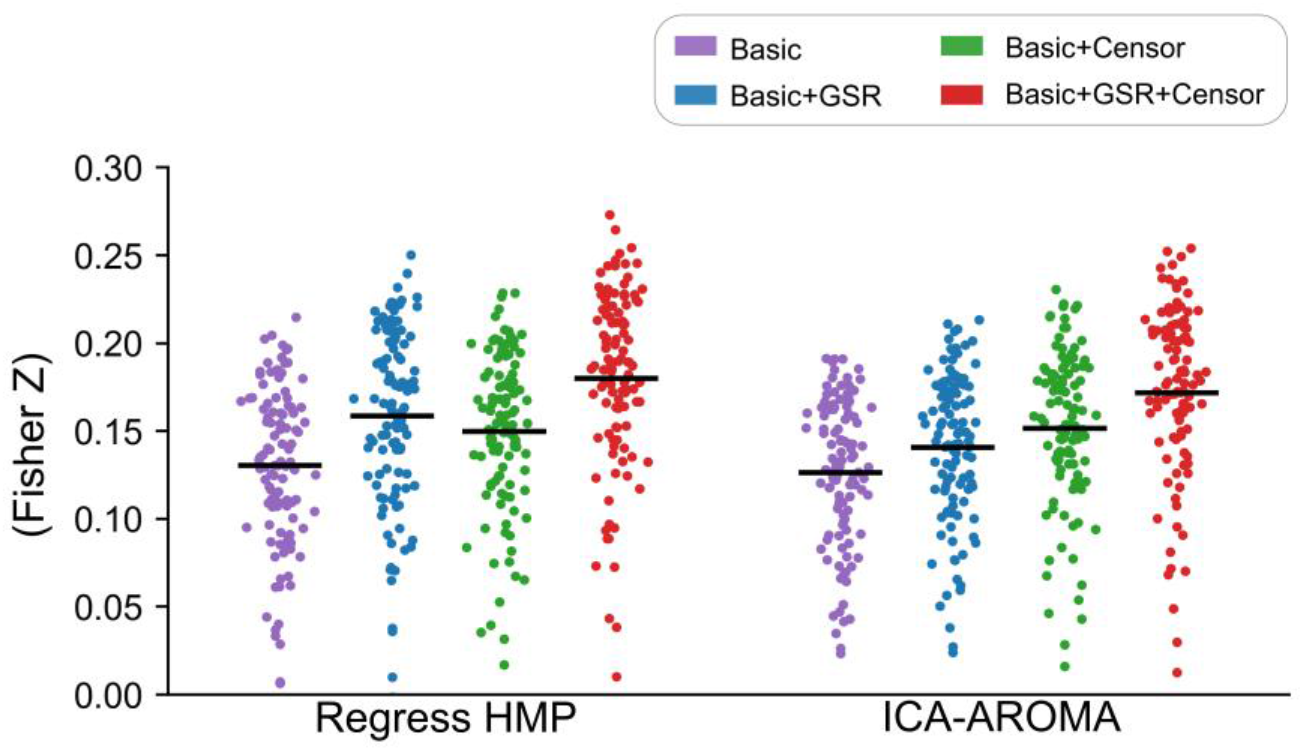
Mean intersubject correlation (ISC) values with a highpass filter. Each point represents the mean ISC for a scan to all other scans, averaged across the 10 parcels with highest ISC. Lines represent mean values. Plot is arranged from pipeline A1 through A8, as described in Table 1.

## References

Aguirre, G. K., Zarahn, E., D’Esposito, M., 1998. The Inferential Impact of Global Signal Covariates in Functional Neuroimaging Analyses. Neuroimage 8, 302–306. https://doi.org/10.1006/nimg.1998.0367

Avants, B.B., Tustison, N.J., Song, G., Cook, P.A., Klein, A., Gee, J.C., 2011. A reproducible evaluation of ANTs similarity metric performance in brain image registration. Neuroimage 54, 2033–2044. https://doi.org/10.1016/j.neuroimage.2010.09.025

Behzadi, Y., Restom, K., Liau, J., Liu, T.T., 2007. A component based noise correction method (CompCor) for BOLD and perfusion based fMRI. Neuroimage 37, 90–101. https://doi.org/10.1016/j.neuroimage.2007.04.042

Bolton, T.A.W., Kebets, V., Glerean., E., Zoller, D., Li., J., Yeo., B.T.T., Caballero-Gaudes., C., Van De Ville., D., 2020. *Agito ergo sum:* Correlates of spatio-temporal motion characteristics during fMRI. Neuroimage 209, 116433. https://doi.org/10.1016/j.neuroimage.2019.116433

Bray, S., Arnold, A.E.G.F., Levy, R.M., Iaria, G., 2015. Spatial and temporal functional connectivity changes between resting and attentive states. Hum. Brain Mapp. 36, 549–565. https://doi.org/10.1002/hbm.22646

Byrge, L., Kennedy, D.P., 2019. High-accuracy individual identification using a “thin slice” of the functional connectome. Netw. Neurosci. 3, 363. https://doi.org/10.1162/NETN_A_00068

Byrge, L., Kennedy, D.P., 2020. Accurate prediction of individual subject identity and task, but not autism diagnosis, from functional connectomes. Hum. Brain Mapp. 41, 2249–2262. https://doi.org/10.1002/hbm.24943

Carp, J., 2012. The secret lives of experiments: Methods reporting in the fMRI literature. Neuroimage 63, 289–300. https://doi.org/10.1016/j.neuroimage.2012.07.004

Chai, X.J., Castañán, A.N., Öngür, D., Whitfield-Gabrieli, S., 2012. Anticorrelations in resting state networks without global signal regression. Neuroimage 59, 1420–1428. https://doi.org/10.1016/j.neuroimage.2011.08.048

Churchill, N.W., Raamana, P., Spring, R., Strother, S.C., 2017. Optimizing fMRI preprocessing pipelines for block-design tasks as a function of age. Neuroimage 154, 240–254. https://doi.org/10.1016/j.neuroimage.2017.02.028

Ciric, R., Wolf, D.H., Power, J.D., Roalf, D.R., Baum, G.L., Ruparel, K., Shinohara, R.T., Elliott, M.A., Eickhoff, S.B., Davatzikos, C., Gur, R.C., Gur, R.E., Bassett, D.S., Satterthwaite, T.D., 2017. Benchmarking of participant-level confound regression strategies for the control of motion artifact in studies of functional connectivity. Neuroimage 154, 174–187. https://doi.org/10.1016/j.neuroimage.2017.03.020

Ciric, R., Rosen, A.F.G., Erus, G., Cieslak, M., Adebimpe, A., Cook, P.A., Bassett, D.S., Davatzikos, C., Wolf, D.H., Satterthwaite, T.D., 2018. Mitigating head motion artifact in functional connectivity MRI. Nat. Protoc. 13, 2801–2826. https://doi.org/10.1038/s41596-018-0065-y

Cox, R.W., 1996. AFNI: Software for analysis and visualization of functional magnetic resonance neuroimages. Comput. Biomed. Res. 29, 162–173. https://doi.org/10.1006/cbmr.1996.0014

Damoiseaux, J.S., Rombouts, S.A.R.B., Barkhof, F., Scheltens, P., Stam, C.J., Smith, S.M., Beckmann, C.F., 2006. Consistent resting-state networks across healthy subjects. Proc. Natl. Acad. Sci. U. S. A. 103, 13848–13853. https://doi.org/10.1073/pnas.0601417103

Davey, C.E., Grayden, D.B., Egan, G.F., Johnston, L.A., 2013. Filtering induces correlation in fMRI resting state data. Neuroimage 64, 728–740. https://doi.org/10.1016/j.neuroimage.2012.08.022

Dimond, D., Rohr, C.S., Smith, R.E., Dhollander, T., Cho, I., Lebel, C., Dewey, D., Connelly, A., Bray, S., 2020. Early childhood development of white matter fiber density and morphology. Neuroimage 210. https://doi.org/10.1016/j.neuroimage.2020.116552

Dimond, D., Heo, S., Ip, A., Rohr, C.S., Tansey, R., Graff, K., Dhollander, T., Smith, R.E., Lebel, C., Dewey, D., Connelly, A., Bray, S., 2020. Maturation and interhemispheric asymmetry in neurite density and orientation dispersion in early childhood. Neuroimage 221. https://doi.org/10.1016/j.neuroimage.2020.117168

Dosenbach, N.U.F., Koller, J.M., Earl, E.A., Miranda-Dominguez, O., Klein, R.L., Van, A.N., Snyder, A.Z., Nagel, B.J., Nigg, J.T., Nguyen, A.L., Wesevich, V., Greene, D.J., Fair, D.A., 2017. Real-time motion analytics during brain MRI improve data quality and reduce costs. Neuroimage 161, 80–93. https://doi.org/10.1016/j.neuroimage.2017.08.025

Fair, D.A., Nigg, J.T., Iyer, S., Bathula, D., Mills, K.L., Dosenbach, N.U.F., Schlaggar, B.L., Mennes, M., Gutman, D., Bangaru, S., Buitelaar, J.K., Dickstein, D.P., Martino, A. Di, Kennedy, D.N., Kelly, C., Luna, B., Schweitzer, J.B., Velanova, K., Wang, Y.F., Mostofsky, S., Castellanos, F.X., Milham, M.P., 2013. Distinct neural signatures detected for ADHD subtypes after controlling for micro-movements in resting state functional connectivity MRI data. Front. Syst. Neurosci. 6, 1–31. https://doi.org/10.3389/fnsys.2012.00080

Finn, E.S., Shen, X., Scheinost, D., Rosenberg, M.D., Huang, J., Chun, M.M., Papademetris, X., Constable, R.T., 2015. Functional connectome fingerprinting: identifying individuals using patterns of brain connectivity. Nat. Neurosci. 18, 1664–71. https://doi.org/10.1038/nn.4135

Fleming, S., Thompson, M., Stevens, R., Heneghan, C., Plüddemann, A., Maconochie, I., Tarassenko, L., Mant, D., 2011. Normal ranges of heart rate and respiratory rate in children from birth to 18 years of age: a systematic review of observational studies. Lancet 377, 1011–1018. https://doi.org/10.1016/S0140-6736(10)62226-X

Fox, M.D., Snyder, A.Z., Vincent, J.L., Corbetta, M., Van Essen, D.C., Raichle, M.E., 2005. From The Cover: The human brain is intrinsically organized into dynamic, anticorrelated functional networks. Proc. Natl. Acad. Sci. 102, 9673–9678. https://doi.org/10.1073/pnas.0504136102

Fox, M.D., Zhang, D., Snyder, A.Z., Raichle, M.E., 2009. The global signal and observed anticorrelated resting state brain networks. J. Neurophysiol. 101, 3270–3283. https://doi.org/10.1152/jn.90777.2008

Friston, K.J., Williams, S., Howard, R., Frackowiak, R.S., Turner, R., 1996. Movement-related effects in fMRI time-series. Magn. Reson. Med. 35, 346–55.

Gordon, E.M., Laumann, T.O., Gilmore, A.W., Newbold, D.J., Greene, D.J., Berg, J.J., Ortega, M., Hoyt-Drazen, C., Gratton, C., Sun, H., Hampton, J.M., Coalson, R.S., Nguyen, A.L., McDermott, K.B., Shimony, J.S., Snyder, A.Z., Schlaggar, B.L., Petersen, S.E., Nelson, S.M., Dosenbach, N.U.F., 2017. Precision Functional Mapping of Individual Human Brains. Neuron 95, 791–807.e7. https://doi.org/10.1016/j.neuron.2017.07.011

Gorgolewski, K., Burns, C.D., Madison, C., Clark, D., Halchenko, Y.O., Waskom, M.L., Ghosh, S.S., 2011. Nipype: A Flexible, Lightweight and Extensible Neuroimaging Data Processing Framework in Python. Front. Neuroinform. 5, 13. https://doi.org/10.3389/fninf.2011.00013

Gotts, S.J., Saad, Z.S., Jo, H.J., Wallace, G.L., Cox, R.W., Martin, A., 2013. The perils of global signal regression for group comparisons: A case study of Autism Spectrum Disorders. Front. Hum. Neurosci. 7. https://doi.org/10.3389/fnhum.2013.00356

Grayson, D.S., Fair, D.A., 2017. Development of large-scale functional networks from birth to adulthood: A guide to the neuroimaging literature. Neuroimage 160, 15–31. https://doi.org/10.1016/j.neuroimage.2017.01.079

Greene, D.J., Koller, J.M., Hampton, J.M., Wesevich, V., Van, A.N., Nguyen, A.L., Hoyt, C.R., McIntyre, L., Earl, E.A., Klein, R.L., Shimony, J.S., Petersen, S.E., Schlaggar, B.L., Fair, D.A., Dosenbach, N.U.F., 2018. Behavioral interventions for reducing head motion during MRI scans in children. Neuroimage 171, 234–245. https://doi.org/10.1016/J.NEUROIMAGE.2018.01.023

Hasson, U., Nir, Y., Levy, I., Fuhrmann, G., Malach, R., 2004. Intersubject Synchronization of Cortical Activity during Natural Vision. Science (80-.). 303, 1634–1640. https://doi.org/10.1126/science.1089506

Huang, C.-M., Lee, S.-H., Hsiao, I.-T., Kuan, W.-C., Wai, Y.-Y., Ko, H.-J., Wan, Y.-L., Hsu, Y.-Y., Liu, H.-L., 2010. Study-specific EPI template improves group analysis in functional MRI of young and older adults. J. Neurosci. Methods 189, 257–66. https://doi.org/10.1016/j.jneumeth.2010.03.021

Jalbrzikowski, M., Liu, F., Foran, W., Klei, L., Calabro, F.J., Roeder, K., Devlin, B., Luna, B., 2020. Functional connectome fingerprinting accuracy in youths and adults is similar when examined on the same day and 1.5-years apart. Hum. Brain Mapp. 41, 4187–4199. https://doi.org/10.1002/hbm.25118

Jenkinson, M., Bannister, P., Brady, M., Smith, S., 2002. Improved optimization for the robust and accurate linear registration and motion correction of brain images. Neuroimage 17, 825–41.

Jo, H.J., Saad, Z.S., Simmons, W.K., Milbury, L.A., Cox, R.W., 2010. Mapping sources of correlation in resting state FMRI, with artifact detection and removal. Neuroimage 52, 571–582. https://doi.org/10.1016/j.neuroimage.2010.04.246

Kaufmann, T., Alnæs, D., Doan, N.T., Brandt, C.L., Andreassen, O.A., Westlye, L.T., 2017. Delayed stabilization and individualization in connectome development are related to psychiatric disorders. Nat. Neurosci. 20, 513–515. https://doi.org/10.1038/nn.4511

Kauppi, J.P., Jääskeläinen, I.P., Sams, M., Tohka, J., 2010. Inter-subject correlation of brain hemodynamic responses during watching a movie: Localization in space and frequency. Front. Neuroinform. 4. https://doi.org/10.3389/fninf.2010.00005

Li, J., Bolt, T., Bzdok, D., Nomi, J.S., Yeo, B.T.T., Spreng, R.N., Uddin, L.Q., 2019. Topography and behavioral relevance of the global signal in the human brain. Sci. Rep. 9, 14286. https://doi.org/10.1038/s41598-019-50750-8

Lindquist, M.A., Geuter, S., Wager, T.D., Caffo, B.S., 2019. Modular preprocessing pipelines can reintroduce artifacts into fMRI data. Hum. Brain Mapp. 40, 2358–2376. https://doi.org/10.1002/hbm.24528

Miranda-Dominguez, O., Feczko, E., Grayson, D.S., Walum, H., Nigg, J.T., Fair, D.A., 2018. Heritability of the human connectome: A connectotyping study. Netw. Neurosci. 02, 175–199. https://doi.org/10.1162/netn_a_00029

Murphy, K., Fox, M.D., 2017. Towards a consensus regarding global signal regression for resting state functional connectivity MRI. Neuroimage 154, 169–173. https://doi.org/10.1016/j.neuroimage.2016.11.052

Niazy, R.K., Xie, J., Miller, K., Beckmann, C.F., Smith, S.M., 2011. Spectral characteristics of resting state networks, in: Progress in Brain Research. pp. 259–276. https://doi.org/10.1016/B978-0-444-53839-0.00017-X

Parkes, L., Fulcher, B., Yücel, M., Fornito, A., 2018. An evaluation of the efficacy, reliability, and sensitivity of motion correction strategies for resting-state functional MRI. Neuroimage 171, 415–436. https://doi.org/10.1016/j.neuroimage.2017.12.073

Power, J.D., Barnes, K.A., Snyder, A.Z., Schlaggar, B.L., Petersen, S.E., 2012. Spurious but systematic correlations in functional connectivity MRI networks arise from subject motion. Neuroimage 59, 2142–54. https://doi.org/10.1016/j.neuroimage.2011.10.018

Power, J.D., Mitra, A., Laumann, T.O., Snyder, A.Z., Schlaggar, B.L., Petersen, S.E., 2014. Methods to detect, characterize, and remove motion artifact in resting state fMRI. Neuroimage 84, 320–341. https://doi.org/10.1016/j.neuroimage.2013.08.048

Power, J.D., Schlaggar, B.L., Petersen, S.E., 2015. Recent progress and outstanding issues in motion correction in resting state fMRI. Neuroimage 105, 536–51. https://doi.org/10.1016/j.neuroimage.2014.10.044

Power, J.D., Plitt, M., Kundu, P., Bandettini, P.A., Martin, A., 2017. Temporal interpolation alters motion in fMRI scans: Magnitudes and consequences for artifact detection. PLoS One 12. https://doi.org/10.1371/journal.pone.0182939

Pruim, R.H.R., Mennes, M., van Rooij, D., Llera, A., Buitelaar, J.K., Beckmann, C.F., 2015. ICA-AROMA: A robust ICA-based strategy for removing motion artifacts from fMRI data. Neuroimage 112, 267–277. https://doi.org/10.1016/j.neuroimage.2015.02.064

Rohr, C.S., Vinette, S.A., Parsons, K.A.L., Cho, I.Y.K., Dimond, D., Benischek, A., Lebel, C., Dewey, D., Bray, S., 2017. Functional Connectivity of the Dorsal Attention Network Predicts Selective Attention in 4–7 year-old Girls. Cereb. Cortex 27, 4350–4360. https://doi.org/10.1093/cercor/bhw236

Rohr, C.S., Dimond, D., Schuetze, M., Cho, I.Y.K., Lichtenstein-Vidne, L., Okon-Singer, H., Dewey, D., Bray, S., 2019. Girls’ attentive traits associate with cerebellar to dorsal attention and default mode network connectivity. Neuropsychologia 127, 84–92. https://doi.org/10.1016/j.neuropsychologia.2019.02.011

Saad, Z.S., Gotts, S.J., Murphy, K., Chen, G., jo, H.J., Martin, A., Cox, R.W., 2012. Trouble at Rest: How Correlation Patterns and Group Differences Become Distorted After Global Signal Regression. Brain Connect. 2, 25–32. https://doi.org/10.1089/brain.2012.0080

Salimi-Khorshidi, G., Douaud, G., Beckmann, C.F., Glasser, M.F., Griffanti, L., Smith, S.M., 2014. Automatic denoising of functional MRI data: Combining independent component analysis and hierarchical fusion of classifiers. Neuroimage 90, 449–468. https://doi.org/10.1016/j.neuroimage.2013.11.046

Satterthwaite, T.D., Wolf, D.H., Loughead, J., Ruparel, K., Elliott, M.A., Hakonarson, H., Gur, R.C., Gur, R.E., 2012. Impact of in-scanner head motion on multiple measures of functional connectivity: relevance for studies of neurodevelopment in youth. Neuroimage 60, 623–32. https://doi.org/10.1016/j.neuroimage.2011.12.063

Satterthwaite, T.D., Elliott, M.A., Gerraty, R.T., Ruparel, K., Loughead, J., Calkins, M.E., Eickhoff, S.B., Hakonarson, H., Gur, R.C., Gur, R.E., Wolf, D.H., 2013. An improved framework for confound regression and filtering for control of motion artifact in the preprocessing of resting-state functional connectivity data. Neuroimage 64, 240–56. https://doi.org/10.1016/j.neuroimage.2012.08.052

Satterthwaite, T.D., Ciric, R., Roalf, D.R., Davatzikos, C., Bassett, D.S., Wolf, D.H., 2019. Motion artifact in studies of functional connectivity: Characteristics and mitigation strategies. Hum. Brain Mapp. 40, 2033–2051. https://doi.org/10.1002/hbm.23665

Smith, S.M., 2002. Fast robust automated brain extraction. Hum. Brain Mapp. 17, 143–155. https://doi.org/10.1002/hbm.10062

Smith, S.M., Jenkinson, M., Woolrich, M.W., Beckmann, C.F., Behrens, T.E.J., Johansen-Berg, H., Bannister, P.R., De Luca, M., Drobnjak, I., Flitney, D.E., Niazy, R.K., Saunders, J., Vickers, J., Zhang, Y., De Stefano, N., Brady, J.M., Matthews, P.M., 2004. Advances in functional and structural MR image analysis and implementation as FSL, in: NeuroImage. Neuroimage. https://doi.org/10.1016/j.neuroimage.2004.07.051

Thomas, C.G., Harshman, R.A., Menon, R.S., 2002. Noise reduction in BOLD-based fMRI using component analysis. Neuroimage 17, 1521–1537. https://doi.org/10.1006/nimg.2002.1200

Urchs, S., Armoza, J., Moreau, C., Benhajali, Y., St-Aubin, J., Orban, P., Bellec, P., 2019. MIST: A multi-resolution parcellation of functional brain networks. MNI Open Res. 1, 3. https://doi.org/10.12688/mniopenres.12767.2

Van Dijk, K.R.A., Sabuncu, M.R., Buckner, R.L., 2012. The influence of head motion on intrinsic functional connectivity MRI. Neuroimage 59, 431–8. https://doi.org/10.1016/j.neuroimage.2011.07.044

Vanderwal, T., Eilbott, J., Finn, E.S., Craddock, R.C., Turnbull, A., Castellanos, F.X., 2017. Individual differences in functional connectivity during naturalistic viewing conditions. Neuroimage 157, 521–530. https://doi.org/10.1016/j.neuroimage.2017.06.027

Vanderwal, T., Eilbott, J., Castellanos, F.X., 2019. Movies in the magnet: Naturalistic paradigms in developmental functional neuroimaging. Developmental Cognitive Neuroscience 36, 100600. https://doi.org/10.1016/j.dcn.2018.10.004

Wang, J., Ren, Y., Hu, X., Nguyen, V.T., Guo, L., Han, J., Guo, C.C., 2017. Test–retest reliability of functional connectivity networks during naturalistic fMRI paradigms. Hum. Brain Mapp. 38, 2226–2241. https://doi.org/10.1002/hbm.23517

